# On-demand delivery of fibulin-1 enables repeated basement membrane stretching

**DOI:** 10.1101/2025.02.02.636120

**Authors:** Adam W.J. Soh, Michael R. Arnwine, Claire A. Gianakas, Zachary D. Clark, Qiuyi Chi, Erin J. Cram, Brenton D. Hoffman, David R. Sherwood

## Abstract

Basement membrane (BM) extracellular matrices enwrap and structurally support tissues. Whether BMs are uniquely constructed to support tissues that undergo repetitive stretching and recoil events is unknown. During *C. elegans* ovulation, the spermathecal BM stretches ∼1.7-fold and then returns to its original shape every twenty minutes to passage hundreds of oocytes. Through live fluorescence microscopy, we discovered that ovulating oocytes secrete and deliver the fibulin-1 extracellular matrix protein to the spermathecal BM during stretching, where it forms a dynamic overlapping network with type IV collagen. Fibulin-1 depletion led to a breakdown in type IV collagen and BM organization, resulting in a more deformable BM and extended spermatheca. Moreover, perturbation to fibulin-1 network formation via mutagenesis was sufficient to disrupt organ shape. Together, our study reveals an on-demand fibulin-1 delivery system that protects the BM network when it is stretched, thereby allowing repeated rounds of organ expansion and recovery.

## Introduction

Tissues such as lung, tendon, and vasculature must repetitively stretch and then return to their original shape, an ability termed elasticity, to serve crucial physiological functions^1^. In vertebrates, elastic fibers and fibrillar collagens work in concert as an elastic system in the connective tissue extracellular matrix (ECM) to support these mechanically active tissues^2^. Elastic fibers, composed primarily of elastin and fibrillin microfibrils, undergo reversible stretching and recoil events in response to tissue deformations, while the fibrillar collagens, largely type I and type III, limit excessive tissue stretching and prevent permanent deformation^3–6^. Highlighting the importance of this elastic system, during aging, elastic fibers fragment and fibrillar collagens accumulate and become disorganized in the lung and aorta, which coincides with reduced elastic recoil (enables recovery to original shape), alveolar enlargement and aorta widening, and reduced tissue function^7–10^.

Basement membranes (BMs) are thin, dense, specialized ECMs built on a scaffolding of type IV collagen and laminin that enwrap most tissues^11, 12^. While the interstitial elastic and collagen fibers can associate with and mechanically support tissue BMs^13–16^, how BMs themselves are constructed to enable reversible deformations is not understood. Inflation and deflation experiments on BM surrounding breast cancer spheroids have revealed that the spheroid BM is elastic (allows expansion and return to its original shape) and has non-linear mechanical properties (increases stiffness at greater stress), which are likely important in maintaining tissue structure^17^. Although the molecular underpinning of BM elasticity remains poorly understood, it has been proposed that the network organization formed by BM proteins might contribute to its ability to reversibly stretch and recoil^17, 18^. However, the identity of BM components and how they are organized in the BM network of mechanically active tissues remain poorly established. The highly abundant and covalently cross-linked type IV collagen is organized as a network that limits tissue deformation^12, 19–21^. Consistent with a potential role for the type IV collagen network in supporting BMs of tissues that are mechanically active, type IV collagen genetic loss leads to embryonic lethality at the onset of muscle contraction in mice, *C. elegans,* and *Drosophila*^22–24^ and mutations in human type IV collagen are associated with defects in mechanically active tissues, such as the vasculature, muscle, and skin^25^. BMs are enormously complex, with over 200 components in vertebrates, and the composition of BMs on each tissue is thought to be tuned for specific signaling and structural functions^26^. The misregulation or genetic alteration of many BM components are also associated with defects in mechanically active tissues, including skin, lung, muscle, and vasculature^26–30^. These observations suggest that BMs of mechanically active tissues might be uniquely enriched with specific matrix proteins that help mediate repetitive BM stretching and recoil events.

Fibulin-1 is a matricellular protein that regulates numerous biological processes, such as embryonic heart and valve development, neural crest cell migration, kidney morphogenesis, and the maturation and organization of elastic fibers in connective tissue^31^. Depletion of fibulin-1 in mice leads to a spectrum of disorders including cranial nerve malformations, thymic hypoplasia, expanded lung sacculi, thinning of cardiac ventricles, and the dilation and rupture of small blood vessels, which likely accounts for the death of most homozygous animals at birth^31, 32^. Fibulin-1 is comprised of three protein domains: domain I harbors three anaphylatoxin-like modules, domain II consists of a series of nine calcium binding EGF-like repeats, and domain III includes a fibulin type module^33^. Electron microscopy and biochemistry studies revealed that fibulin-1 forms a 20 nm dumbbell-shaped rod that self-associates laterally via EGF repeats 5 and 6 in a calcium-dependent manner^34, 35^. Electron microscopy imaging has further revealed that fibulin-1 associates through end-to-end on interactions, although the terminus mediating these interactions is unclear^35^. Fibulin-1 also binds numerous BM components, such as type IV collagen, laminin, nidogen, and proteoglycans^35, 36^. The role of fibulin-1 in supporting the BM of dynamic tissues in vivo, however, is unclear, in large part because of the challenge of visualizing and experimentally examining BM component regulation and function in mechanically active tissues that repeatedly expand and recover their shape in native settings.

Here, we develop *Caenorhabditis elegans* ovulation as an in vivo experimental model to examine how BM supports elastic tissues that repetitively stretch and return to their original shape. Using live imaging, photobleaching and volumetric analysis, we show that the BM associated with the spermathecal organ repeatedly stretches ∼1.7 fold then reversibly recovers during the hundreds of ovulation events that occur for fertilization and passage of zygotes to the uterus. Through a visual screen of over 50 endogenous mNeonGreen (mNG) tagged BM proteins, we identified fibulin-1 as uniquely enriched in the spermathecal BM in a type IV collagen dependent manner. Conditional knockdown studies revealed that loss of fibulin-1 initially had little effect on BM stretching and ovulation. However, after repeated ovulations, the integrity of the type IV collagen and laminin networks was disrupted and the spermathecal BM became more deformable when stretched (the BM overstretched), coinciding with a loss of spermathecal shape and function. Using live imaging and targeted oocyte expression, we discovered that oocytes deliver fibulin-1 to the spermathecal BM specifically during ovulation when it is stretched. Our imaging studies revealed that type IV collagen and fibulin-1 assemble into overlapping networks that dynamically expand when the BM stretches. Perturbation to the fibulin-1 network was sufficient to lead to the formation of extended spermathecas, highlighting the importance of network organization on organ shape. Together, these results reveal a dynamic delivery mechanism that supplies fibulin-1 to the spermathecal BM when it is stretched, where it forms into a network that protects the BM, thus enabling repeated cycles of organ extension and return to original shape.

## Results

### The *C. elegans* spermathecal basement membrane (BM) stretches during ovulation

*C. elegans* have two bilaterally symmetric gonad arms that connect at a central uterus through opposing spermathecal organs (Fig. 1a). At the proximal ends of the gonad, mature oocytes are driven individually into the spermatheca by contractions of the gonadal sheath cells. Upon entry to the spermatheca, an oocyte is fertilized by stored sperm and through actomyosin dependent contractions of the spermathecal myoepithelial cells, the zygote is expelled from the spermatheca and enters the uterus^37–40^. To determine how the spermathecal BM that enwraps the organ stretches and retracts during ovulation, we first wanted to verify that the BM surrounding the spermatheca is in continual contact with the spermathecal organ during ovulation and establish the degree and time it takes the spermatheca to expand and recover to its resting shape (Fig. 1). Self-fertile hermaphrodites begin ovulating during day 1 of adulthood at a rate of ∼40 progeny per day and reach peak progeny production of ∼120 on day 2. Ovulation decreases to ∼60 progeny per day on day 3 and ceases by day 5^41^. Using timelapse confocal fluorescence microscopy of day 1 and 2 adult animals, along with semi-automated volumetric analysis (Methods), we found that the spermathecal BM (visualized by the endogenously tagged core BM laminin, LAM-2::mNG^42^) continually enwrapped the spermathecal tissue (visualized by spermatheca specific expression of the F-actin probe, mCherry::moeABD^43^) during ovulation (Fig.1b,c and Supplementary Video 1). We also discovered that the spermatheca expanded 2-fold in volume (Fig. 1d and Extended Data Fig. 1a), and there was 1.5-fold increase in length and 3-fold increase in circumference (Fig. 1c,e,f). Oocyte entry into the spermatheca took approximately 3 min, full spermathecal expansion lasted for ∼2 min, and then retraction occurred over a 3 min period, which returned the spermatheca to the resting size (Fig. 1b-d and Supplementary Video 1). These observations demonstrate that during ovulation the BM is in tight association with the spermathecal organ, which dramatically expands and returns to its resting shape.

**Figure 1.**
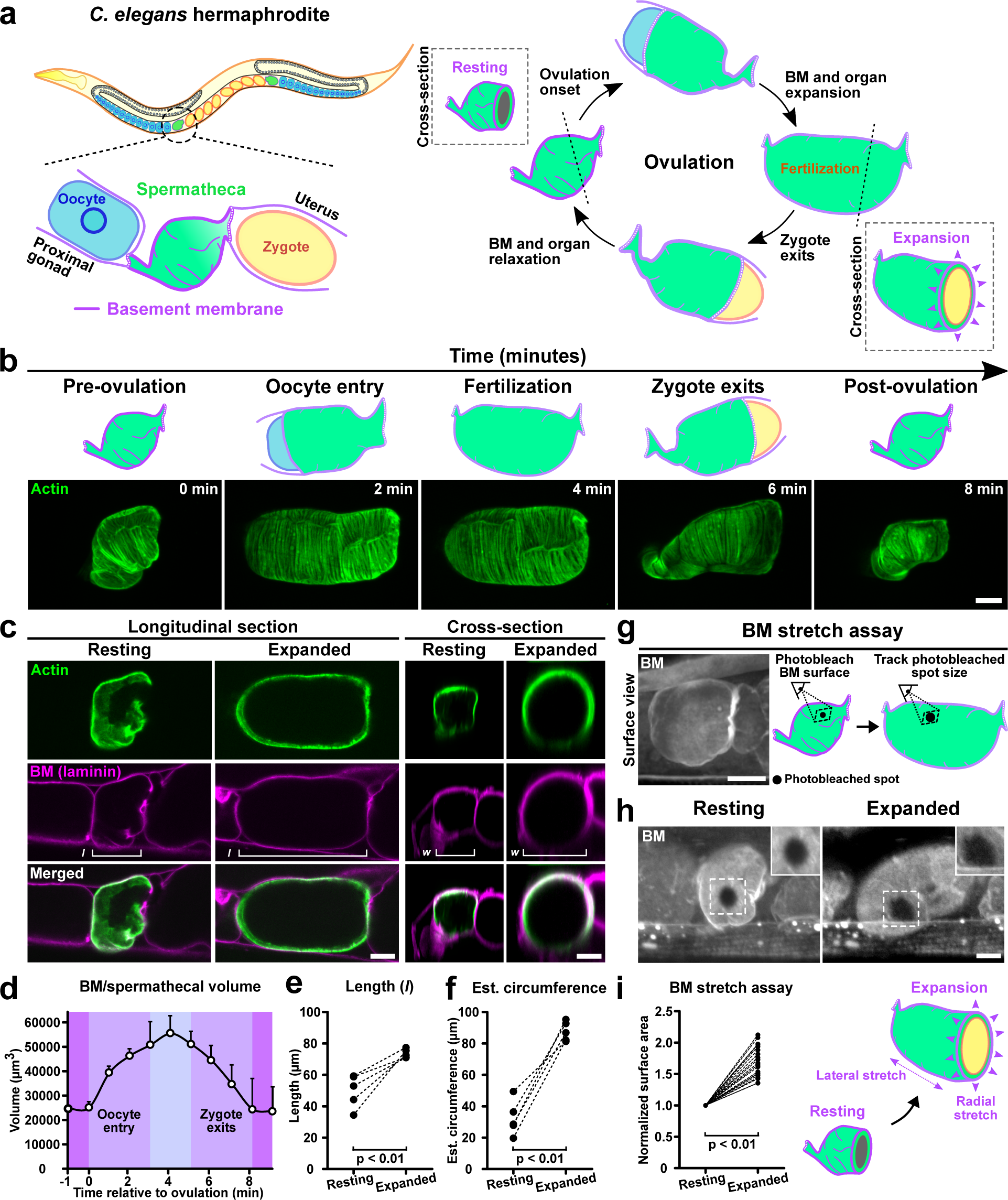
The *C. elegans* spermathecal basement membrane (BM) stretches during ovulation. **a**, A schematic diagram of the *C. elegans* gonad in an adult hermaphrodite showing spermathecal expansion and return to the resting state during ovulation. **b,** Maximum intensity z-projected fluorescence timelapse images showing the spermatheca (actin, mCherry::moeABD, green) during ovulation. **c,** Longitudinal and cross-sectional views of the spermatheca (mCherry::moeABD, green) and its enveloping BM (laminin γ1 chain, LAM-2::mNG, magenta). *l*, length and *w*, width. **d,** Quantification of the BM/spermathecal volume during ovulation (n = 5 animals; Mean + SD). **e and f,** Quantification of the spermathecal length and circumference during the resting and expanded states (n = 5 animals; *Mann-Whitney* test). **g,** (*Left*) Maximum intensity z-projected fluorescence image showing the surface of the spermathecal BM (LAM-2::mNG, gray) during the resting state. (*Right*) A schematic diagram of the spermathecal BM stretch assay. **h,** Maximum intensity z-projected fluorescence images of the spermathecal BM surface (type IV collagen α2 chain, LET-2::mRuby, gray) showing a photobleached spot in the resting and expanded state. Inset shows a magnified view of the photobleached spot (inset width is 12 µm). **i,** Quantification of the normalized surface area of photobleached spots in resting and stretched BMs (n = 20 animals; Student’s *t-*test). All data is representative of 3 independent biological repeats. Scale bars, 10 µm.

The core BM components of laminin and type IV collagen stably associate with BM in the gonadal and pharyngeal tissues^42^. To directly test if the BM stretches, a 5 µm diameter circle of fluorescent type IV collagen (LET-2::mRuby) was photobleached in a region devoid of BM folds and the bleached region was followed during ovulation (Fig. 1g; Methods). The photobleached spot increased ∼70% in size at full spermathecal expansion during ovulation (Fig. 1h,i). Furthermore, the BM stretched more along the circumferential axis than the longitudinal axis of the spermatheca, mirroring spermathecal organ expansion (Fig. 1e,f and Extended Data Fig. 1b). To independently test these observations, we also photobleached a 5 µm diameter circle in fluorescent laminin (LAM-2::mNG) and observed similar BM stretching (Extended Data Fig. 1c). Finally, we examined the fluorescence intensity of spermathecal BM laminin and type IV collagen during ovulation. The fluorescence intensity of type IV collagen and laminin decreased by ∼50% upon BM stretching (Extended Data Fig. 3a,b), consistent with stretching of the spermathecal BM and thus reduction in the density of fluorescently tagged components. Together, our findings indicate that the spermathecal BM stretches when the spermathecal organ expands to accommodate oocytes during ovulation.

### The spermathecal BM has a unique composition

We next wanted to determine if the spermathecal BM has a unique composition that might facilitate stretching. We previously identified 98 genes encoding BM associated proteins in *C*. *elegans*, which includes 71 matrix components and 27 cell surface interactors^26^. We have endogenously tagged 55 of these BM components with mNeonGreen (mNG) that are homozygous viable. These include all core BM matrix components, most BM associated receptors and many matricellular proteins, enzymes, and proteases^26, 42^. These strains allow localization and quantitative comparisons of BM component levels. We performed a visual screen of all tagged BM proteins and identified 16 clearly localized to the spermathecal BM (Fig. 2a-c, Extended Data Fig.1d,e and Supplementary Table 1). Another nine BM proteins were in the extracellular fluid surrounding the spermatheca or within the spermathecal myoepithelial cells, and did not appear to concentrate in the spermathecal BM (Extended Data Fig.1d and Supplementary Table 1). Spermathecal BM localized components include the core BM proteins type IV collagen (LET-2 and EMB-9 α chains), papilin (MIG-6), nidogen (NID-1), type XVIII collagen (CLE-1), laminin (EPI-1 α, LAM-1 β, and LAM-2 γ chains) and perlecan (UNC-52). We also identified the BM accessory matrix components SPOCK (TEST-1), SPARC (OST-1), peroxidasin-1 (PXN-1), and fibulin-1 (FBL-1), as well as the cell surface integrin heterodimer receptor (INA-1/PAT-3), dystroglycan receptor (DGN-1), UNC-5 and DCC netrin 1 receptors (UNC-5 and UNC-40), ROBO receptor (SAX-3) and the PTPRD receptor (PTP-3) (Fig. 2b,c and Extended Data Fig. 1d). Quantitative fluorescence analysis revealed that 10 out of the 16 components were enriched at the spermathecal BM compared with the adjoining proximal gonad and uterine BMs (Fig. 2b,c and Extended Data Fig. 1e and 2). Notably, type IV collagen levels (LET-2) were ∼3 fold higher and nidogen ∼2 fold higher in the spermathecal BM, while laminin levels (LAM-2) were modestly higher (∼1.2-fold; Fig. 2c and Extended Data Fig. 2).

**Figure 2.**
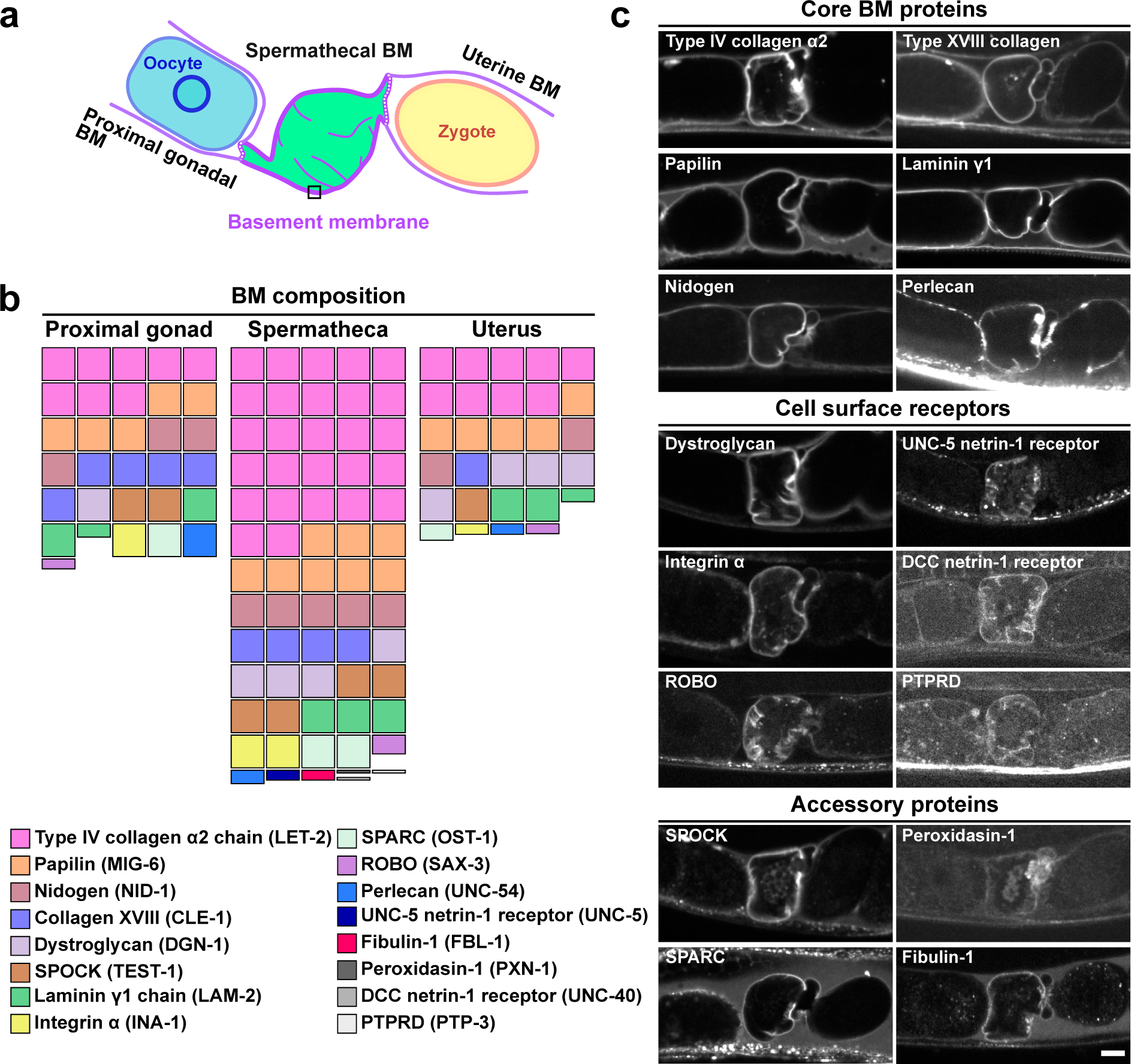
The spermathecal BM has a distinct composition. **a**, A schematic diagram showing the proximal gonad (teal), resting spermatheca (green), and uterus enwrapped by a continuous BM (purple). The fluorescence intensity of each BM protein was sampled at the proximal gonadal BM, spermathecal BM (box), and uterine BM.**b,** Normalized fluorescence intensity of BM protein abundance displayed as waffle plots to compare the composition of the proximal gonadal BM, spermathecal BM, and uterine BM. All fluorescence intensities were normalized to the average fluorescence intensity of LAM-2::mNG (laminin γ1 chain) in resting spermathecal BMs (n = 30-50 animals for all BM proteins, except for LAM-2::mNG (n = 145 animals)). **c,** Fluorescence images (single z-slice) of BM proteins that localize to the spermathecal BM. Scale bar, 10 µm.

Comparison of the spermathecal BM composition between resting and stretched BMs revealed that the composition was ∼50% lower in fluorescence density in the stretched state, adding further evidence that the entire BM scaffolding expands during ovulation (Extended Data Fig. 3a,b). We conclude that the spermathecal BM composition is distinct from the adjoining proximal gonadal and uterine BMs.

### The spermathecal BM and organ shape depends on specific spermathecal BM components

Specific BM proteins, such as type IV collagen, laminin, perlecan, and proteases shape and support the integrity of tissues during their formation in development^44–50^. Whether specific BM proteins maintain the shape and function of organs that repeatedly stretch and return to their original shape is unclear. The *C. elegans* spermatheca consists of 24 myoepithelial cells that are arranged to form a flexible tube that fully develops by the late larval L4 stage^38, 51^ (Fig. 3a). To determine if BM components are required to support the integrity of the spermathecal organ as it undergoes repeated cycles of expansion and recovery, we performed a conditional RNAi screen using feeding RNAi beginning at the L4 stage when the spermatheca has formed and then assessed spermathecal shape in day 1 and 2 adults (24 h and 48 h later) after the onset of ovulation. We screened BM proteins that localize to the spermathecal BM as well as BM proteins that were found in the vicinity of the spermathecal BM (Fig. 2b,c and Extended Data Fig. 1d). Depletion of the core BM components of type IV collagen, laminin and papilin led to high penetrance of elongated spermatheca (>85%) near the onset of ovulation (Extended Data Fig. 3c and Supplementary Table 2), suggesting that these components are required for the spermathecal shape at the onset of ovulation. An elongated spermatheca was also observed at lower penetrance (47%) after nidogen depletion, and the penetrance increased modestly (63%) after a day of ovulation (Supplementary Table 2). The depletion of the remaining two core BM components, type XVIII collagen and perlecan, did not affect spermathecal shape nor did reduction of the cell surface receptors, proteases, and the accessory components SPARC, SPOCK and the enzymes peroxidasin-1 and peroxidasin-2 (Supplementary Table 2). Depletion of fibulin-1, a matricellular BM component, resulted in a low penetrance of elongated spermatheca (30%) at the onset of ovulation. Notably, the penetrance of the spermathecal shape defect increased dramatically to ∼95% during peak ovulation (Supplementary Table 2). To rule out the possibility that fibulin-1 was not depleted at the earlier time point, fibulin-1 (*fbl-1*) RNAi knockdown efficiency was assessed and confirmed to be comparable at both times (Extended Data Fig. 4a,b). These results suggest that the core BM components of type IV collagen, papilin (promotes type IV collagen turnover^42^), and laminin are required within the BM to maintain spermathecal shape from the onset of ovulation, but that fibulin-1 might have a specific role in maintaining the spermathecal BMs ability to undergo repeated stretching and recovery events and support organ shape.

**Figure 3.**
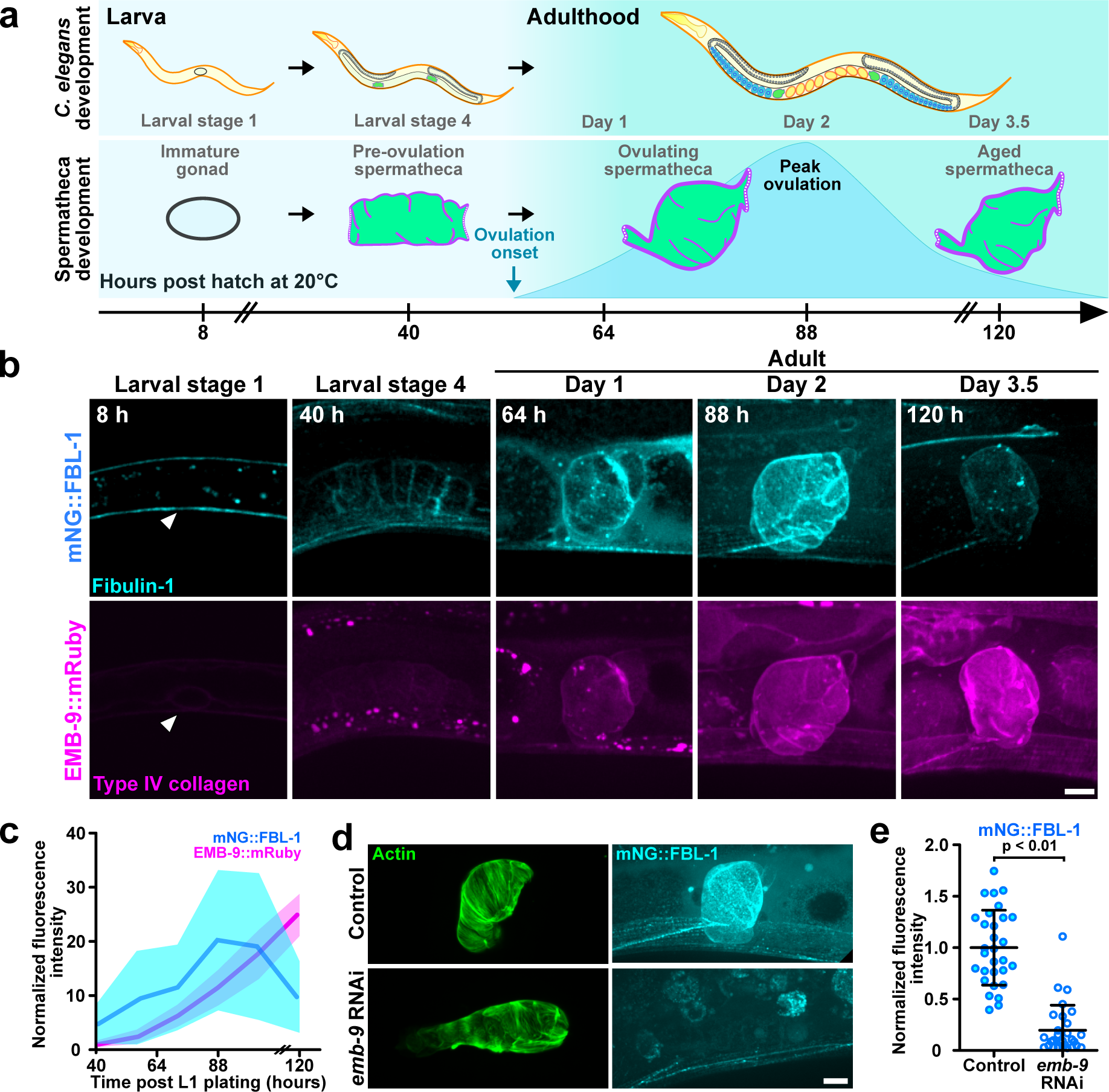
Fibulin-1 requires type IV collagen for localization and enrichment in the spermathecal BM during peak ovulation. **a**, A schematic diagram showing the timing of *C. elegans* spermathecal development. The spermatheca is formed during the late L4 stage. Ovulation begins shortly after the L4 stage, and peak ovulation occurs in day 2 adult animals. The ovulation rate dramatically declines in day 3.5 adult animals. **b,** Maximum intensity z-projected fluorescence images showing the levels of fibulin-1 (mNG::FBL-1, cyan) and type IV collagen α1 chain (EMB-9::mRuby, magenta) from the late L4 stage to day 3.5 adult animals. Fibulin-1 was not detectable in the gonadal BM during the L1 stage (arrowhead). **c,** Quantification of fibulin-1 and type IV collagen levels in the spermathecal BM (n = 28-32 animals per time point; *Mann-Whitney* test; mean ± SD). **d,** Maximum intensity z-projected fluorescence images showing the spermathecal BM fibulin-1 (cyan) and organ (actin, mCherry::moeABD, green) in control and type IV collagen depleted (*emb-9* RNAi) day 2 adult animals. **e,** Quantification of fibulin-1 levels in control and type IV collagen depleted (*emb-9* RNAi) day 2 adult animals (control, n = 30 animals; RNAi, n = 29 animals; *Mann-Whitney* test; mean ± SD). All data is representative of 3 independent biological repeats. Scale bars, 10 µm.

### Fibulin-1 is most enriched in the spermathecal BM during peak ovulation and requires type IV collagen for localization

To begin to understand how fibulin-1 sustains the integrity of the spermatheca during repeated ovulations, we first determined when fibulin-1 (mNG::FBL-1) is incorporated into the spermathecal BM. Fibulin-1 was not detectable in the gonadal BM of L1 stage animals and only became detectable in the spermathecal BM when the organ first forms at the late L4 stage (40h, (Fig. 3a,b). Fibulin-1 levels, however, dramatically increased at the initiation of ovulation with the highest levels occurring in day 2 adult animals at the peak of ovulation (Fig. 3b,c). Fibulin-1 levels then decreased in day 3.5 adult animals when ovulation events diminished (Fig. 3b,c). Unlike fibulin-1, the levels of the core BM component type IV collagen (EMB-9:: mRuby) continued to increase during the same time window (Fig. 3b,c).

Biochemical studies have shown that vertebrate fibulin-1 binds to type IV collagen and laminin, which are required for proper spermathecal shape from the onset of ovulation^35^. We were thus interested in determining if these proteins mediate fibulin-1 recruitment to BM. Using feeding RNAi beginning at the L4 stage, we found that depletion of type IV collagen reduced fibulin-1 levels dramatically (∼80%) at the spermathecal BM in day 2 adult animals (Fig. 3d,e), whereas RNAi mediated reduction of laminin did not impact fibulin-1 levels (Extended Data Fig. 4c,d). While papilin is not known to interact with fibulin-1, it is required for proper spermathecal shape (Supplementary Table 2). Depletion of papilin, however, did not affect fibulin-1 levels in the spermathecal BM (Extended Data Fig. 4c,d). We conclude that increased fibulin-1 levels in the spermathecal BM are correlated with increased ovulation frequency and require type IV collagen for BM enrichment.

### Fibulin-1 maintains spermathecal shape and ovulation function

To investigate the role of fibulin-1 in maintaining organ shape, we next examined how loss of fibulin-1 affects the shape of the spermatheca at different times over the course of ovulation— onset (Day 1 adult), early (Day 1.5 adult) and peak ovulation (Day 2 adult). *fbl-1* RNAi was initiated at the L4 stage just prior to the onset of ovulation, which reduced fibulin-1 level by at least ∼80% throughout ovulation (Fig. 4a and Extended Data Fig. 4a,b). Organ shape was assessed by measuring the length of the spermatheca during the resting state (when the oocyte is absent) between ovulations (Fig. 4b,c). We found at the onset of ovulation that the overall length of the BM enwrapped spermatheca (BM/spermatheca) of fibulin-1 depleted animals was slightly (but not significantly) longer (Day 1 adult; Fig. 4d), consistent with our initial screen identifying a small number of animals with extended spermathecas at this time (Supplementary Table 2).

**Figure 4.**
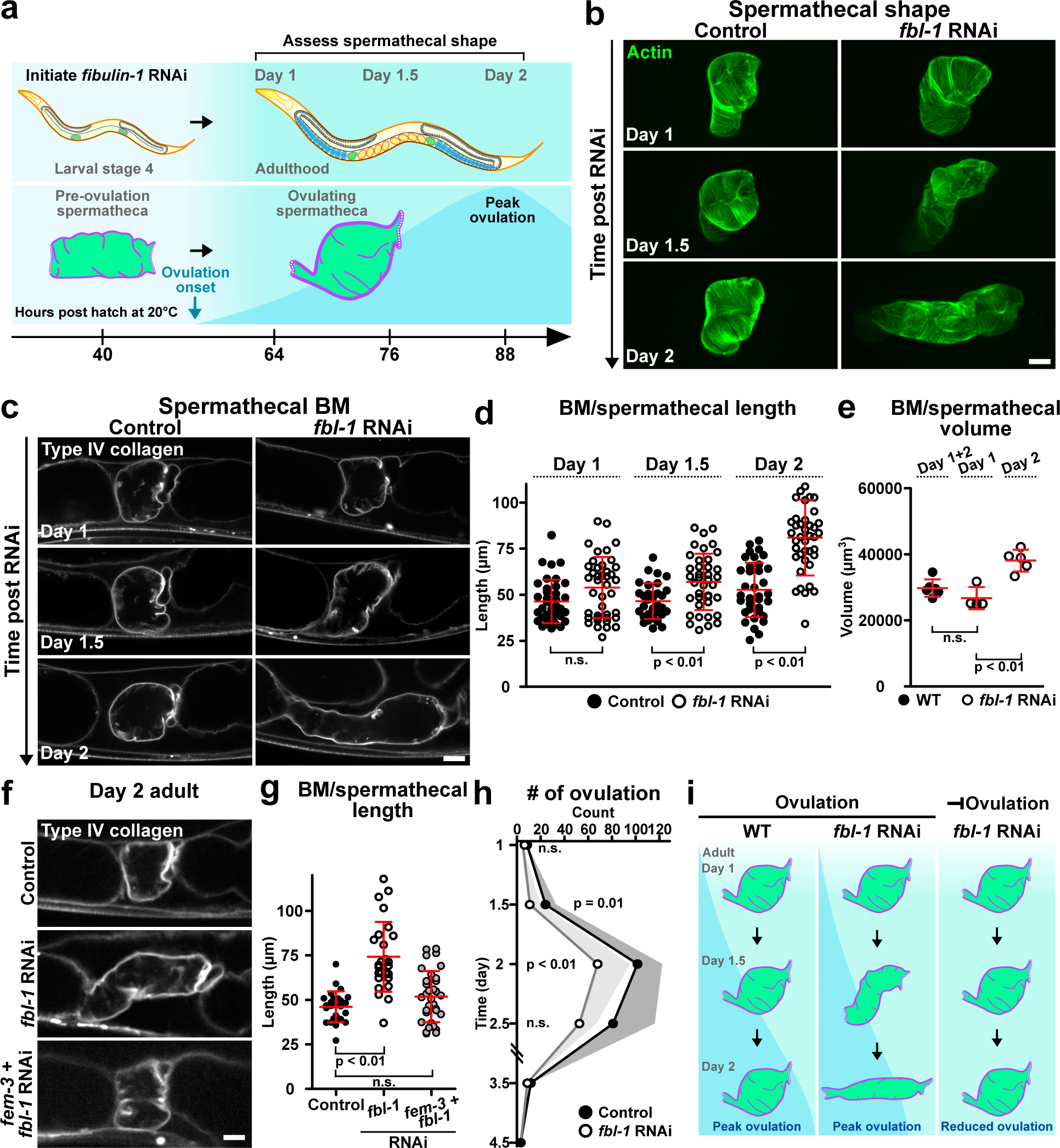
Fibulin-1 maintains spermathecal shape and function after repeated rounds of ovulations. **a**, A schematic diagram describing the assay to study how fibulin-1 supports spermathecal shape. **b,** Maximum intensity z-projected fluorescence images showing the spermathecal organ (actin, mCherry::moeABD , green) in control and *fbl-1* RNAi treated day 1-2 adult animals. **c,** Fluorescence images (single z-slice) showing the spermathecal BM (type IV collagen α2 chain, LET-2::mRuby, gray) in control and *fbl-1* RNAi treated day 1-2 adult animals. **d,** Quantification of spermathecal length in control and *fbl-1* RNAi treated day 1-2 adult animals (n = 31-41 animals per time point; *Mann-Whitney* test; mean ± SD). **e,** Quantification of spermathecal volume in control and *fbl-1* RNAi treated day 1-2 adult animals (control in day 1-2 adult animals, n = 6 animals; *fbl-1* RNAi in day 1 adult animals, n = 4 animals; *fbl-1* RNAi in day 2 adult animals, n = 5 animals; *Mann-Whitney* test; mean ± SD). **f,** Fluorescence images (single z-slice) showing the spermathecal BM (LET-2::mRuby, gray) in control, *fbl-1* RNAi, and combinatorial (*fem-3* and *fbl-1*) RNAi treated day 2 adult animals. **g,** Quantification of spermathecal length in control, *fbl-1* RNAi, combinatorial (*fem-3* and *fbl-1*) RNAi treated day 2 adult animals (n = 26-33 animals; Student’s *t*-test; mean ± SD). **h,** Quantification showing the number of ovulation events in control and *fbl-1* RNAi treated day 1-4.5 adult animals (n = 10 animals; *Mann-Whitney* test; mean + SD). **i,** A schematic diagram summarizing the role of fibulin-1 in sustaining spermathecal shape after repeated ovulation events. All data is representative of 3 independent biological repeats. Scale bars, 10 µm.

The spermatheca continued to elongate at the early stages of ovulation (Day 1.5 adult) and lengthened dramatically during peak ovulation (Day 2 adult; Fig. 4b-d). The BM/spermathecal volume in fibulin-1 depleted animals was also comparable to control animals at the onset of ovulation (Day 1 adult; Fig. 4e), but the volume increased by ∼28% during peak ovulation (Day 2 adult; Fig. 4b,e), confirming the progressive elongation of the spermatheca after fibulin-1 loss.

To determine if repeated ovulations drive progressive spermatheca extension after fibulin-1 loss, we next assessed if elongated spermathecas arose in the absence of ovulation. To inhibit ovulation, worms were co-treated with *fem-3* and *fbl-1* RNAi. The FEM-3 protein blocks sperm production, which is required for oocyte maturation and ovulation^52, 53^. We found that in non-ovulating animals lacking fibulin-1, the spermatheca was comparable in length relative to control animals during the usual time of peak ovulation (Fig. 4f,g). These results strongly support the notion that fibulin-1 is required to maintain spermathecal organ shape as it undergoes repeated rounds of stretching and recovering during ovulation.

We next examined ovulation events at different time points to determine if loss of fibulin-1 affected ovulation (Methods). We found that fibulin-1 depleted (*fbl-1* RNAi) animals had a normal number of ovulations events at the onset of ovulation, but the number of ovulations decreased by ∼30% during the time of peak ovulation in day 2 adults (Fig. 4h). Timelapse analysis further revealed ovulation was unaffected initially, but between day 1.5-2, ∼30% of oocytes did not exit the spermatheca for the duration of the timelapse (up to 80 min) (Extended Data Fig. 5a-c and Supplementary Video 2). The expulsion of the zygote from the spermatheca into the uterus is driven by coordinated contraction of the actomyosin network in the spermathecal myoepithelial cells and this process is dependent on the organization of the actin cytoskeleton^38, 54^. Unlike the closely positioned actin bundles that were observed in both control and fibulin-1 depleted animals at the onset of ovulation during the resting state, we found that distance between spermathecal actin bundles increased by ∼40% at the time of peak ovulation in fibulin-1 depleted animals (Extended Data Fig. 5d,e), suggesting that the perturbation of ovulation upon fibulin-1 loss might in part be due to defective actin organization and contractile ability. We conclude that in the absence of fibulin-1 there is a progressive decline in spermathecal shape and function after repeated ovulations.

### Loss of fibulin-1 perturbs spermathecal BM stretching

As fibulin-1 is assembled into the spermathecal BM, we were next interested in understanding if loss of fibulin-1 affected the stretching of the BM during ovulation. We thus performed the photobleaching stretch assay on type IV collagen after RNAi mediated depletion of fibulin-1 protein (Fig. 5a). At the onset of ovulation (Day 1 adult), we found that both control and fibulin-1 depleted animals displayed a comparable extent of BM stretching (Fig. 5a,b). During peak ovulation, however, the spermathecal BM in fibulin-1 depleted animals stretched ∼30% more than control animals (Day 2; Fig. 5a,b). Importantly, the height and width of the zygote in control animals were similar to those in fibulin-1 depleted animals, thus the increase in BM stretching was not attributable to increased zygote size (Fig. 5c). Together, these results suggest that loss of fibulin-1 leads to a more deformable BM after repeated ovulations (Fig. 5d).

**Figure 5.**
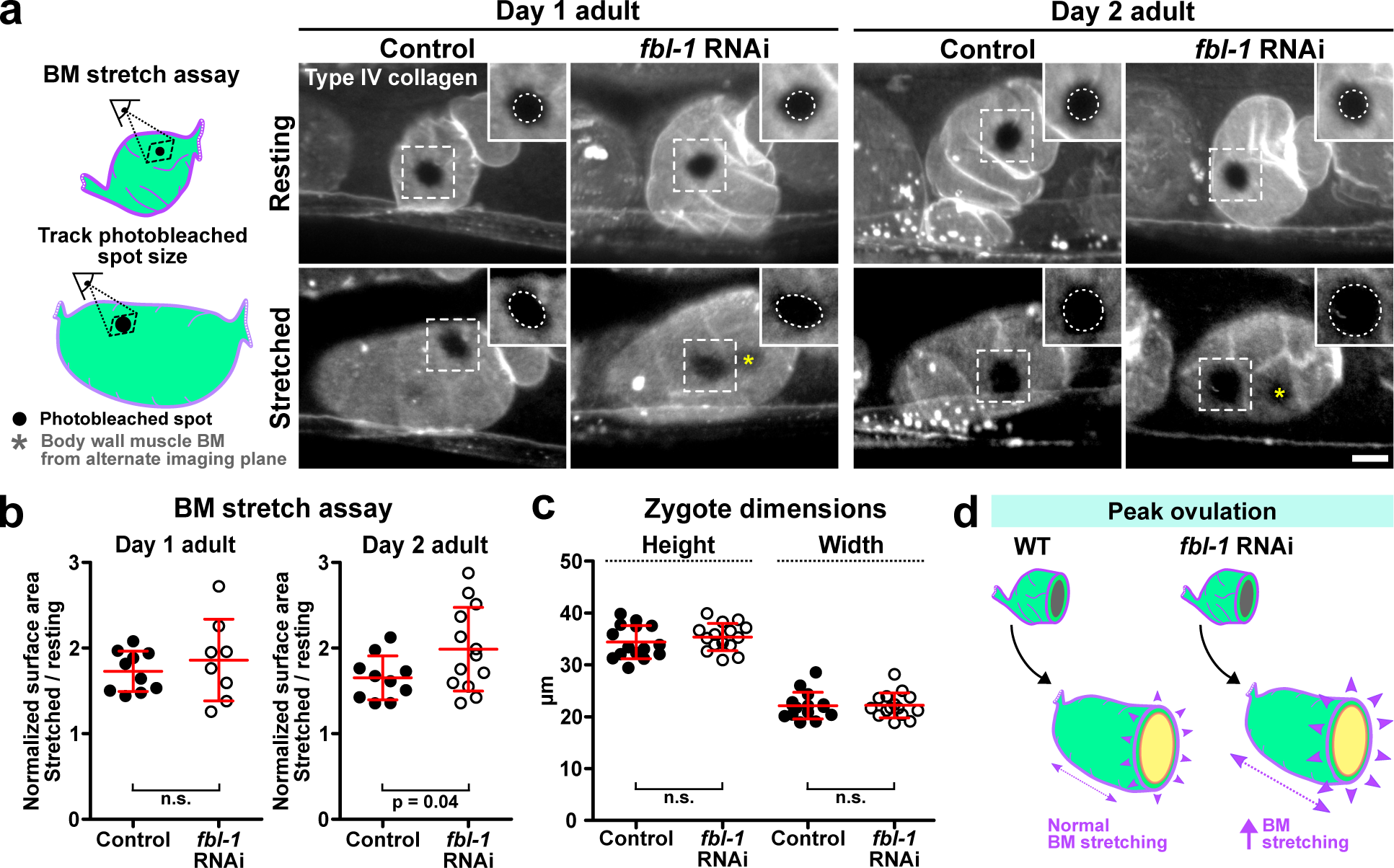
Fibulin-1 loss perturbs spermathecal BM stretching. **a**, (*Left*) A schematic diagram of the spermathecal BM stretch assay. (*Right*) Maximum intensity z-projected fluorescence images showing the surface view of resting and stretched BMs (LET-2::mRuby, gray) in control and *fbl-1* RNAi treated day 1 and 2 adult animals. Inset shows a magnified view of the photobleached spot (inset width is 12 µm). Images showing resting BMs were uniformly contrasted. The brightness of images showing stretched BMs was increased by a factor of 2. Due to its proximity, the body wall muscle BM was sometimes photobleached (yellow asterisk) and projected in the same field of view. **b,** Quantification showing the normalized surface area of photobleached spots in control and *fbl-1* RNAi treated day 1 and 2 adult animals (n = 9-13 animals; Student’s *t*-test with Welch’s correction; mean ± SD). **c,** Quantification of zygote length and width (n =15 animals; Student’s *t*-test; mean ± SD). **d,** A schematic diagram showing that in the absence of fibulin-1, the BM stretches farther than untreated (normal) BMs during peak ovulation. All data is representative of 3 independent biological repeats. Scale bars, 10 µm.

### Fibulin-1 is delivered by oocytes to the spermathecal BM specifically during ovulation

Fibulin-1 levels in the spermathecal BM appeared tuned to ovulation activity, increasing dramatically during peak ovulation and then declining (Fig. 3b). To further examine the link between ovulation and fibulin-1 levels, we used *fem-3* RNAi treatment, which inhibits ovulation, and found fibulin-1 levels were dramatically reduced (decreased ∼50%, Extended Data Fig. 5f,g). To determine how fibulin-1 delivery to the spermathecal BM might be linked to ovulation, we carried out timed analysis of fibulin-1 presence within the spermathecal myoepithelial cells. Fibulin-1, however, was not detected in resting or stretched spermatheca myoepithelial cells (Extended Data Fig. 5h-j). Instead, we observed that the most proximal oocyte was enriched in fibulin-1 puncta in ovulating animals (Fig. 6a). Timelapse analysis revealed that fibulin-1 became detectable in the oocyte ∼30 minutes prior to ovulation and then dramatically increased, reaching peak levels just prior spermatheca entry, and then rapidly decreased as the oocyte passed through the spermatheca, which coincided with increased fibulin-1 levels in the extracellular fluid surrounding the spermathecal BM (Fig. 6a-c and Supplementary Video 3). FRAP experiments revealed that ∼40% of fibulin-1 fluorescence signal within the resting BM recovered within 10 minutes after photobleaching indicating that fibulin-1 has rapid on-off dynamics on the BM (Extended Data Fig. 6a,b and Supplementary Video 4). FRAP experiments on type IV collagen did not show significant fluorescence recovery during the same experimental time window (Extended Data Fig. 6c,d and Supplementary Video 4). These results suggest that ovulating oocytes secrete fibulin-1, which dynamically associates with the spermathecal BM.

**Figure 6.**
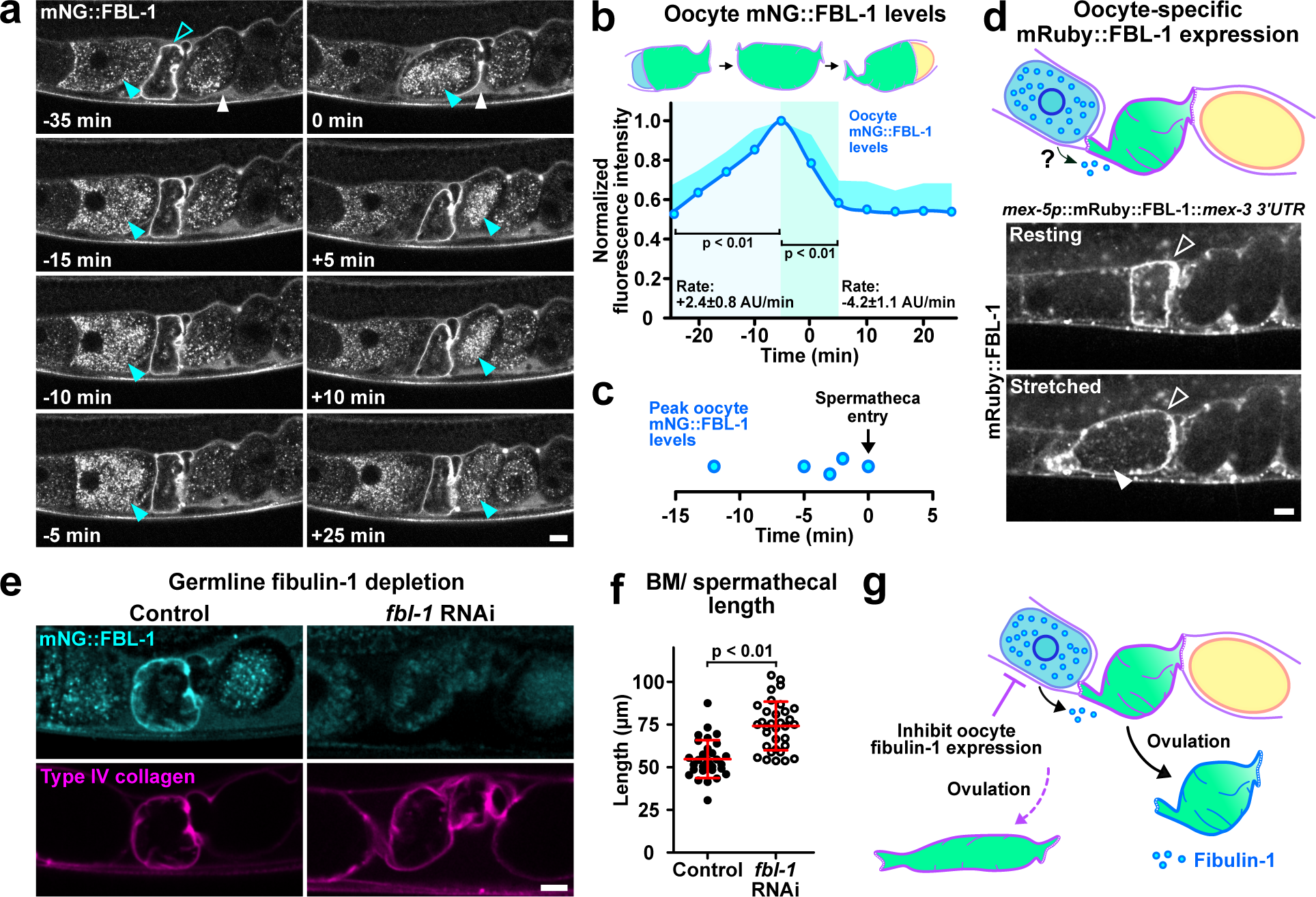
Ovulating oocytes provide fibulin-1 to the spermathecal BM. **a**, Fluorescence timelapse images (single z-slice) showing that fibulin-1 (mNG::FBL-1, gray) is present and builds in levels within vesicles (cyan arrowhead) in a representative oocyte just before it enters the spermatheca, is present in vesicles and the BM (open arrowhead) of the spermatheca, and then fibulin-1 levels decline in the vesicles of zygote after it leaves the spermatheca (cyan arrowhead). Fibulin-1 is also at high levels in the extracellular fluid (white arrowhead) surrounding the spermatheca (n = 5 animals). **b,** Quantification of oocyte fibulin-1 levels through the course of an ovulation event (n = 5 animals; *Mann-Whitney* test; mean + SD). **c,** Quantification showing that oocyte fibulin-1 level peak prior to entry into the spermatheca (0 min) (n = 5 animals). **d,** (*Top*) A schematic diagram describing the oocyte-specific expression strategy to determine if oocytes secrete and deliver fibulin-1 to the spermathecal BM. (*Bottom*) Fluorescence images (single z-slice) showing that oocyte-expressed fibulin-1 (mRuby::FBL-1) localized to both resting and stretch spermathecal BMs (open arrowhead) and was also detected within the ovulating oocyte (filled arrowhead; n = 10 animals). **e,** Fluorescence images (single z-slice) showing fibulin-1 abundance (mNG::FBL-1, cyan) at the spermathecal BM (LET-2::mRuby, magenta) of control and germline-specific *fbl-1* RNAi treated day 2 adult animals. **f,** Quantification of spermathecal length in control and germline-specific *fbl-1* RNAi treated day 2 adult animals (Control, n = 31; RNAi, n = 33 animals; Student’s *t*-test; mean ± SD). **g,** A schematic diagram showing ovulating oocytes providing fibulin-1 to the spermathecal BM. All data is representative of 3 independent biological repeats. Scale bars, 10 µm.

To assess the contribution of oocyte fibulin-1 to the spermatheca BM, we first determined if oocytes are sufficient to deliver fibulin-1 to the spermathecal BM. We used an oocyte-specific transgene expression strategy^55^ (*mex5p::mRuby::fbl-1::mex3* 3’ UTR) and found that oocyte expressed mRuby::FBL-1 was present in spermathecal BM in day 1-2 adults (Fig. 6d, Extended Data Fig. 6e and Supplementary Video 5). To determine the functional role of fibulin-1 secreted from the oocytes, we next used germline-specific RNAi, which inhibits fibulin-1 expression from the germ cells including the oocytes^56^. Germline depletion of fibulin-1 led to ∼50% reduction in fibulin-1 spermathecal BM levels and this was sufficient to result in elongated spermathecas during peak ovulation (Fig. 6e-g and Extended Data Fig. 6f). Taken together, these results indicate that oocytes deliver fibulin-1 to the spermathecal BM to maintain spermathecal organ shape during ovulation (Fig. 6g).

### Fibulin-1 forms a network within the BM and maintains BM integrity

Fibulin-1 binds type IV collagen, laminin and nidogen^35^. We hypothesized that fibulin-1 could act to maintain spermathecal BM structure during repeated stretching and recovery events. To assess type IV collagen integrity in the spermathecal BM, we first examined the type IV collagen fluorescence signal across the spermathecal BM surface at the onset of ovulation. Both control and fibulin-1 depleted animals displayed uniform distribution of type IV collagen fluorescence signal at the onset of ovulation, suggesting intact type IV collagen integrity (Fig. 7a-c and Extended Data Fig. 7a,b). However, after repeated cycles of ovulation, gaps in the type IV collagen fluorescence signal (average width of 1.8 µm) appeared in ∼60% of fibulin-1 depleted animals but only ∼6% in control animals (Fig. 7a-c and Extended Data Fig. 7a,b), suggesting a breakdown in type IV collagen integrity. We also observed gaps in laminin in ∼90% of fibulin-1 depleted animals by peak ovulation (Extended Data Fig. 7c), suggesting the entire BM was disrupted after multiple ovulation events.

**Figure 7.**
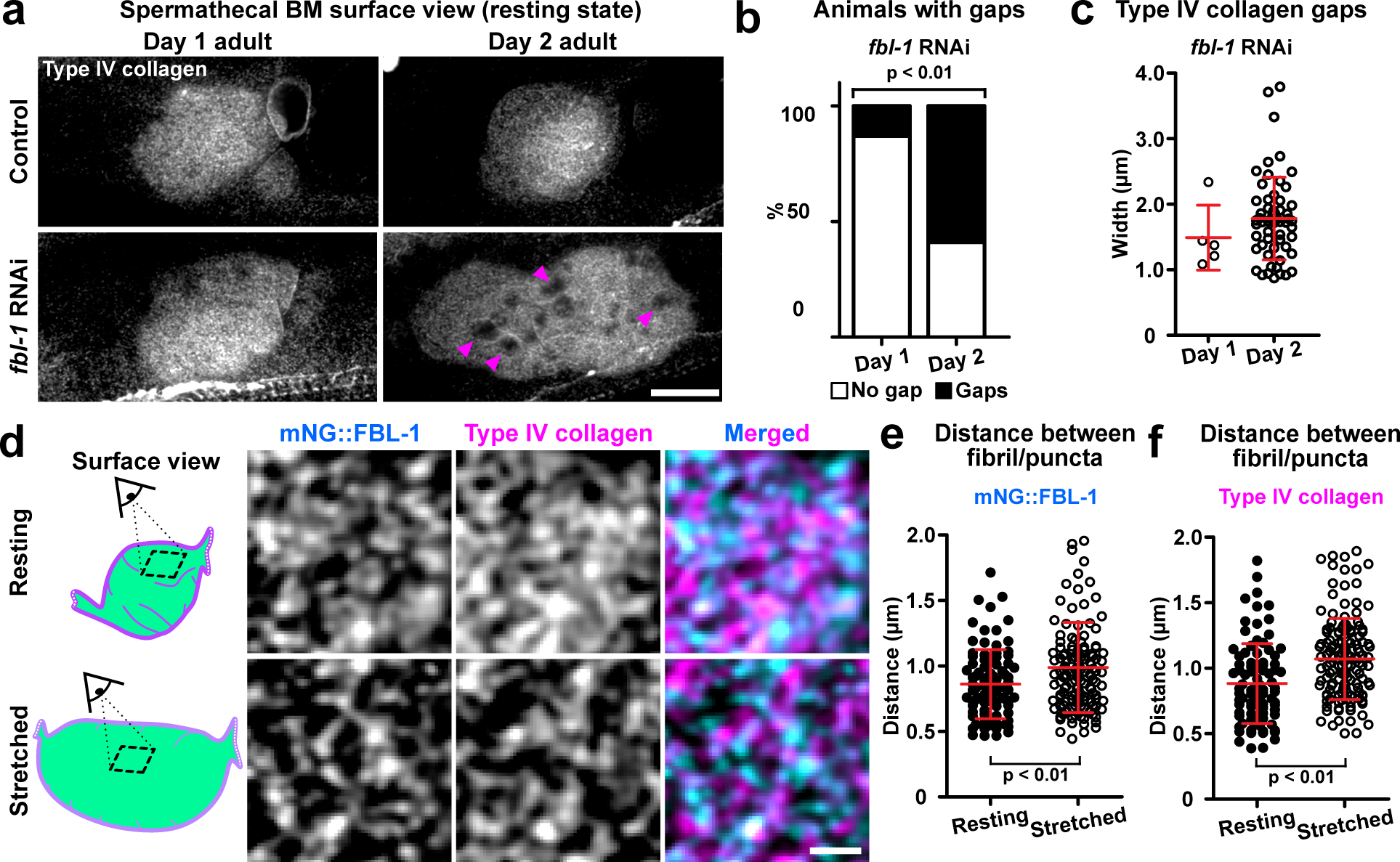
Fibulin-1 forms a network that maintains the type IV collagen network. **a**, Maximum intensity z-projected fluorescence images showing the surface view of the type IV collagen within the spermathecal BM (α2 chain, LET-2::mRuby, gray) in control and *fbl-1* RNAi treated day 1 and 2 adult animals. Gaps (arrowheads) in the type IV collagen network were detected in *fbl-1* RNAi treated day 2 adult animals. **b,** Proportion of animals with type IV collagen gaps in *fbl-1* RNAi treated day 1 and 2 adult animals. (*fbl-1* RNAi day 1 adult animal, n = 22 animals; *fbl-1* RNAi day 2 adult animal, n = 27 animals; *Fisher’s exact* test). **c,** Quantification of type IV collagen gap widths in *fbl-1* RNAi treated day 1 and 2 adult animals (*fbl-1* RNAi day 1 adult animal, n = 22 animals; *fbl-1* RNAi day 2 adult animal, n = 27 animals; *Mann-Whitney* test; mean ± SD). **d,** Fluorescence images (single z-slice) showing fibulin-1 (mNG::FBL-1, cyan) and type IV collagen α2 chain (LET-2::mRuby, magenta) form overlapping networks in the spermathecal BM. Both networks expand when the spermathecal BM stretches. **e,** Quantification of distance between fibulin-1 fibril/puncta in resting and stretched BMs (resting state n = 90 pairs of fibril/puncta from 9 animals; Stretched state n = 140 pairs of fibril/puncta from 14 animals; *Mann-Whitney* test; mean ± SD). **f,** Quantification of distance between type IV collagen fibril/puncta in resting and stretched BMs (resting state n = 90 pairs of fibril/puncta from 9 animals; stretched state n = 140 pairs of fibril/puncta from 14 animals; *Mann-Whitney;* mean ± SD). All data is representative of 3 independent biological repeats. Scale bars, 10 µm (**a**) and 1 µm (**d**).

To further test the hypothesis that fibulin-1 maintains type IV collagen and BM integrity, we assessed if the gaps in the spermathecal type IV collagen fluorescence signal are more likely to form when the spermathecal BM is subjected to increased stretching. To increase BM stretching, animals were treated with filamin-1 (*fln-1*) RNAi, which leads to multiple oocytes being trapped in the spermatheca^57^. Examination of animals with two oocytes trapped in the spermatheca at the start of ovulation, revealed that increased BM stretching led to a higher frequency of animals with gaps in type IV collagen fluorescence signal (average width: ∼1.3 µm; Extended Data Fig. 7d-f). Notably, the co-depletion of filamin-1 and fibulin-1 via combinatorial RNAi treatment led to a 3-fold increase in animals with gaps in the type IV collagen fluorescence signal (Extended Data Fig. 7d-f). Importantly, the levels of type IV collagen, laminin and papilin (core BM components) were similar to control animals after fibulin-1 loss (Extended Data Fig. 7g-m), and nidogen levels were only modestly reduced (∼25%, Extended Data Fig. 7g,n,o). Thus, fibulin-1 maintains type IV collagen and BM integrity in the spermathecal BM.

Isolated fibulin-1 molecules have the capacity to self-associate^34^, suggesting that fibulin-1 might form a network within BM. To test this, we assessed fibulin-1 organization in resting and stretched spermathecal BM using confocal microscopy. We found that fibulin-1 is organized as a heterogenous meshwork of fibrils and puncta across the BM with an average distance between fibril/puncta of ∼0.85 µm in resting spermathecal BMs and ∼1.00 µm in stretched spermathecal BMs (Fig. 7d,e). We next examined how the fibulin-1 network is organized relative to type IV collagen. Like fibulin-1, we found that type IV collagen forms a heterogenous network of fibrils and puncta in the spermathecal BM that expands by ∼20% when the BM was stretched during ovulation (Fig. 7d,f; ∼0.90 µm resting versus ∼1.10 µm stretched). Co-occurrence analysis showed colocalization of fibulin-1 and type IV collagen (∼20%) and distinct meshworks (∼40%) in the resting and stretched states (Fig. 7d and Extended Data Fig. 7p,q). These findings indicate

that fibulin-1 and type IV collagen form overlapping networks that expand when the spermathecal BM stretches.

### Fibulin-1 network maintains spermathecal shape

Given that fibulin-1 forms a meshwork in the spermathecal BM and that fibulin-1 is required to maintain the spermathecal BM and organ shape after repeated ovulations, we next wanted to determine if the network organization of fibulin-1 is required for fibulin-1 function. Biochemistry studies suggest that fibulin-1 monomers might self-associate laterally via EGF repeats 5 and 6 in a calcium dependent manner and by end-on associations through an unknown mechanism^34, 35^. These self-associating interactions might have a role in fibulin-1’s ability to form a network^58^. Thus, we next examined if the inhibition of fibulin-1 lateral associations was sufficient to disrupt network formation. Using site-directed mutagenesis, the cysteine residue in EGF repeat 5 at position 413 was replaced with a phenylalanine residue (mNG::FBL-1(C413F)) to disrupt calcium ion binding^34, 59^ (Fig. 8a). To determine if the fibulin-1 meshwork was disrupted, we exogenously expressed mNG::FBL-1(C413F) and assessed its localization. Importantly, all comparisons were carried out on animals expressing similar levels of mNG::FBL-1 (wild-type) and mNG::FBL-1(C413F) (mutant) (Extended Data Fig. 8a). Unlike wild-type fibulin-1, which was organized into a clear network, the C413F mutant formed puncta that were shorter in length than control animals in both resting and stretched spermathecal BMs (Fig. 8b). Thus, the mNG::FBL-1(C413F) mutant is capable of localizing to the spermathecal BM but shows reduced network forming ability.

**Figure 8.**
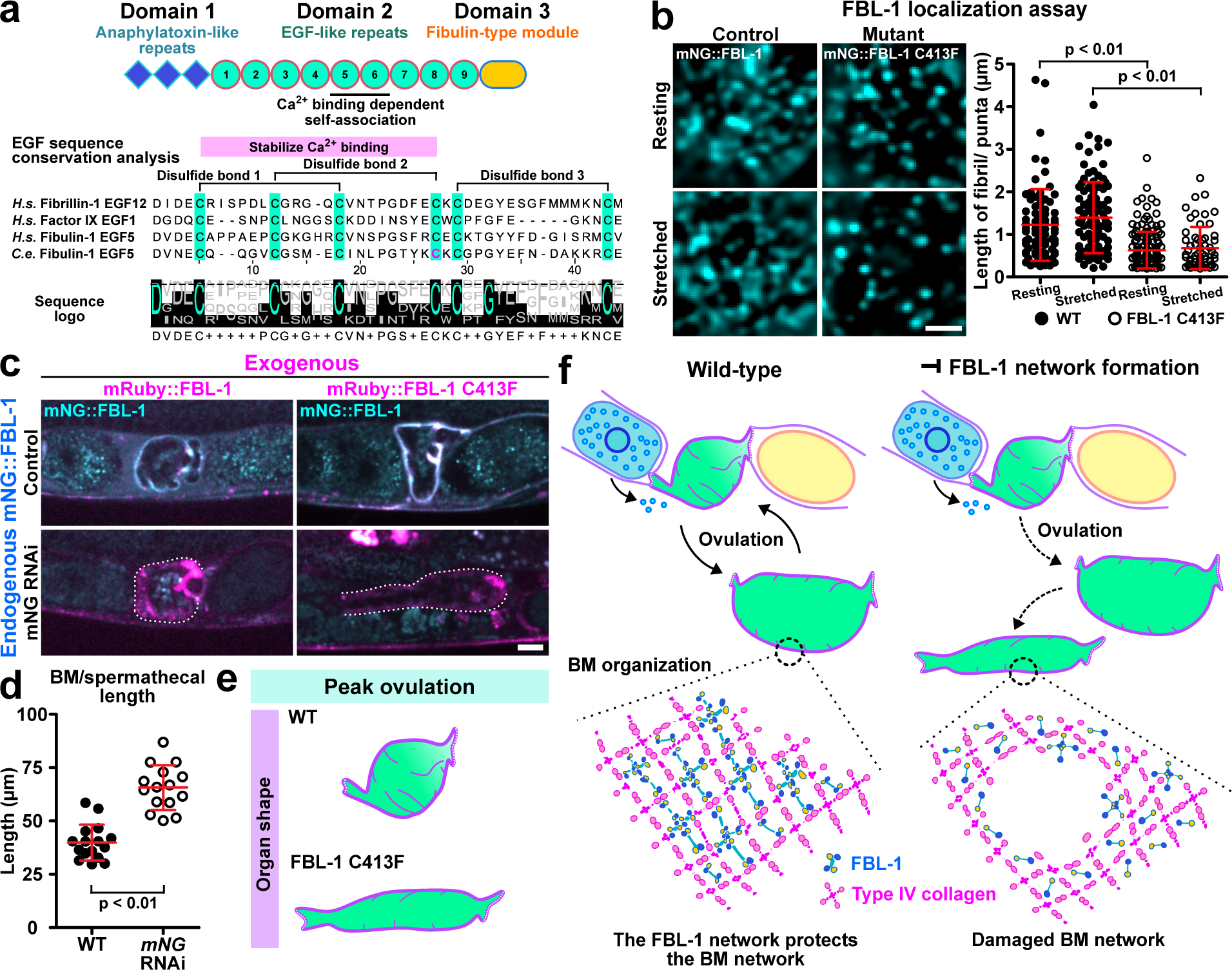
Fibulin-1 network maintains spermathecal shape. **a**, Phylogenetic analysis assessing the conservation of *C. elegans* fibulin-1 EGF repeat 5 relative to the human fibulin-1 EGF repeat 5, factor IX EGF repeat 1 and fibrillin EGF repeat 12. The cysteine (C) residue at amino acid position 413 (magenta) was substituted with a phenylalanine (F) residue to disrupt calcium ion binding and fibulin-1 self-association. **b,** (*Left*) Fluorescence images (single z-slice) showing the localization of wild-type and C413F mutant fibulin-1 (mNG::FBL-1,cyan) at the spermathecal BM. (*Right*) Quantification of WT and C413F mutant fibril/puncta length (wild-type fibulin-1, resting state n = 80 pairs of fibrils from 8 animals; stretched state n = 100 pairs of fibrils from 10 animals; C413F mutant fibulin-1, resting state n = 160 pairs of fibrils from 16 animals; stretched state n = 60 pairs of fibrils from 6 animals; *Mann-Whitney* test; mean ± SD). **c,** Fluorescence images (single z-slice) showing exogenously expressed wild-type or C413F mutant fibulin-1 (mRuby::FBL-1). The mNG tagged endogenous fibulin-1 (mNG::FBL-1) was depleted via *mNG* RNAi. **d,** Quantification of spermathecal length of wild-type fibulin-1 or C413F mutant in day 2 adult animals (wild-type fibulin-1, n = 16 animals; C413F mutant fibulin-1, n = 15 animals; Student’s *t*-test; mean ± SD). **e,** A schematic diagram showing that the C413F mutant fibulin-1 was unable to maintain spermathecal shape. **f,** A schematic diagram summarizing the proposed role of oocyte delivered fibulin-1 in forming a network that maintains the integrity of the spermathecal BM, which supports organ shape and ovulation. All data is representative of 3 independent biological repeats. Scale bar, 1 µm.

We next wanted to determine if the fibulin-1 meshwork is required for proper organ shape. By adapting the conditional RNAi strategy described in Fig. 3a, we depleted the endogenous population of fibulin-1 (mNG::FBL-1) via RNAi targeting mNG and assessed if the spermathecal shape was affected in the presence of either exogenously expressed wild-type mRuby::fibulin-1 or mRuby::C413F mutant. Control animals expressing wild-type mRuby::FBL-1 displayed normal spermathecal length (Fig. 8c,d and Extended Data Fig. 8b), indicating that the exogenous fibulin-1 rescued the depletion of endogenous fibulin-1 and maintained spermathecal shape. Conversely, animals that expressed mRuby::FBL-1 (C413F) displayed extended spermathecas when endogenous fibulin-1 was depleted (Fig. 8c-e and Extended Data Fig. 8b,c). We conclude that the fibulin-1 network is required for fibulin-1’s function in maintaining spermathecal shape (Fig. 8f).

## Discussion

Many tissues must repeatedly stretch and then return to their original shape to function. A major obstacle in elucidating how BMs support these mechanically active tissues has been the lack of in vivo models that allow BMs, BM components, and the tissue that the BM enwraps to be visualized and manipulated. Here, we developed the *C. elegans* spermatheca as a model to study how BMs support tissues that repeatedly expand and return to their resting shape. During ovulation, an oocyte enters the spermatheca, which causes the organ to expand. After fertilization, the embryo is rapidly expelled, and the spermatheca returns to its initial shape. At peak egg laying, ovulation events occur every twenty minutes, and the spermatheca can withstand hundreds of ovulations^41^. Using volumetric analysis and photobleaching, we found that the spermathecal BM stretches ∼1.7 fold during a single ovulation event. Through a visual screen of over 50 endogenously tagged BM components coupled to an RNAi screen, we discovered that fibulin-1 is uniquely present in the spermathecal BM and is required to maintain the shape and function of the spermathecal organ during repeated ovulations. Notably, mice deficient in fibulin-1 have expanded lung sacculi and widened capillaries^32^—defects in the shape of tissues that also undergo cyclic expansion and recovery. Thus, fibulin-1 may have a conserved role within BMs in supporting tissues that repeatedly stretch and recoil.

Our live cell imaging also revealed an elegant delivery on-demand system that provides fibulin-1 to the BM when it is needed during stretching. Fibulin-1 was present as puncta in the cytosol of oocytes prior to ovulation and throughout the time of ovulation, as well as in the extracellular fluid surrounding the spermathecal BM, suggesting that the oocytes secrete fibulin-1 during ovulation to provide it to the spermathecal BM. Supporting this idea, oocyte-specific expression of fibulin-1 was sufficient to deliver fibulin-1 to the spermathecal BM and germline-specific RNAi dramatically decreased fibulin-1 in the spermathecal BM and led to a loss of spermathecal shape. Similar to oocyte delivered fibulin-1 in the lumen of the spermatheca, mammalian fibulin-1 is found at high levels in the blood plasma and increases significantly during placentation when expansion of the vasculature occurs^60–62^. As fibulin-1 is required to support vascular integrity^31, 32^, a high extracellular reservoir might be required to maintain the BMs of expanding and retracting capillaries.

Studies on BMs surrounding breast cancer spheroids using a microneedle to inflate the BM and then follow its behavior, have revealed that spheroid BMs can expand and then return to their initial size, suggesting that BMs are inherently elastic^17^. The mechanisms underlying BMs elasticity are poorly understood but have been suggested to be dependent on the individual proteins and network organization of BMs^17, 18^. Type IV collagen is a core BM component that assembles into a cross-linked network that is crucial to support BMs and tissue structure^20, 21^.

Consistent with the idea that the BM network organization is elastic, our in vivo studies showed that type IV collagen assembles into an irregular meshwork in the spermathecal BM that stretches and then returns to its original state. Our imaging studies also revealed that fibulin-1 forms a network that overlaps with the type IV collagen network and that it expands and contracts during ovulation. Depletion of fibulin-1 led to a breakdown of the type IV collagen network after repeated cycles of stretching and recovery and disrupted the laminin network, suggesting that fibulin-1 protects the BM from damage. We found that fibulin-1 function was dependent on self-association^34^ and network formation. Given that fibulin-1 binds many BM proteins, including type IV collagen, laminin and nidogen^35, 63^, it suggests that the fibulin-1 network might protect the BM by binding to many components of the BM network. The oocyte delivery system for fibulin-1, and rapid on-off dynamics at the spermathecal BM (half-life ∼4 min), further suggests that fibulin-1 is likely added to the BM during stretching and recovery, thus providing an adaptable network that guards the BM from damage.

During ageing, elastic fibers and fibrillar collagens that are aligned along the aorta fragment, become disorganized and collagens accumulate, and this is associated with a loss of the stretching and recovery properties of the aorta^7^. Progressive lengthening and widening of the aorta also occur, indicating a failure of the aorta tissue to return to its resting state^7, 8^. The observation that fibulin-1 depletion also led to a progressive lengthening of the spermatheca suggests that in the absence of fibulin-1 the damaged BM has reduced elastic recoil as there is a failure to recover the BM enwrapped tissue to its original shape. In vitro studies have indicated that BMs can undergo elastic recoil^17^ and the depletion of the core BM components of type IV collagen, laminin and nidogen resulted in extended spermathecas from the onset of ovulation, suggesting that the core BM network proteins contribute to elastic recoil. As type IV collagen contains flexible interruptions within the collagenous domains and form extensible bundles of intertwined collagen fibrils in the network^20, 21, 64^, type IV collagen might directly contribute to the BMs ability to return to its original shape after stretching.

BMs are complex supramolecular networks that impart tissues with unique mechanical properties to resist external and internal forces and serve as signaling hubs throughout an animal’s lifetime^12, 49^. During aging, the BMs of many mechanically active tissues such as the skin and vasculature decline, coinciding with disrupted tissue shape and function^65, 66^. Interesting, fibulin-1 levels decrease during aging, suggesting that its loss might contribute to the decline of BMs^67^. Understanding how fibulin-1 maintains the integrity and elasticity of BMs in the face of recurring mechanical stresses is thus important as it might lead to better diagnosis and treatment strategies for BM-associated human diseases.

## Author contributions

A.S. and D.S. conceived the project and wrote the manuscript. A.S. designed, conducted all experiments and analyses. M.A. assisted with the characterization of spermathecal BM stretching and examined the role of fibulin-1 on spermathecal BM stretching and organ shape. C.G. assisted with the characterization of BM stretching and volume, and conducted a subset of the visual screen. Z.C. conducted a subset of the visual screen to identify spermathecal BM proteins and quantified their relative abundance. Q.C. designed and built molecular biology constructs. E.C. designed the experiment to study the role of fibulin-1 on the spermathecal actin cytoskeleton. A.S., D.S., E.C. and B.H. read and commented on the manuscript.

## Methods

### Strains and culture conditions

*C. elegans* strains were fed *E. coli* (OP50) and cultured on nematode growth media (NGM) plates at 20°C as previously described^68^. Strains used in this study are summarized in Supplementary Table 4. Strains sourced from Caenorhabditis Genetics Center (CGC) are to be requested directly from CGC. Further information and requests for reagents should be directed to the correspondent authors Adam W.J. Soh and David Sherwood.

### Construction of fusion proteins

In this study, the linkage between a promoter (p), open reading frame, and fluorophore is designated with a double semicolon (::)^69^.The germline specific expression of *mRuby::FBL-1* was constructed by PCR fusion to generate the transgene *mex-5p::mRuby::fbl-1::mex-3* 3’ UTR. A 1.9 kbp fragment of the *mex-5* promoter region, full length *fbl-1* genomic sequence, and 0.7 kbp fragment of the *mex-3* 3’ UTR were fused in the pAP088 vector. The expression of mNG::FBL-1 was constructed by PCR fusion to generate the transgene *fbl-1p::mNG::fbl-1::fbl-1* 3’ UTR. A 1.9 kbp fragment of the *fbl-1* promoter region, full length *fbl-1* genomic sequence (5.2 kbp), and 0.7 kbp fragment of the *fbl-1* 3’ UTR were fused in the pAP088 vector. The same promoter and 3’UTR sequences were used for the expression of the mNG::FBL-1 C413F mutant. mRuby tagged versions of wild-type and mutant fibulin-1 were made using the same strategy. The construct and Cas9-sgRNA plasmid (pCFJ352) were co-injected into the gonad of young adult N2 hermaphrodites. Recombinant animals were identified in the F3 offspring of injected animals via selectable markers (hygromycin resistance and dominant-negative *sqt-1 rol* phenotype)^70^.

### Construction of the fibulin-1 Cys413Phe point mutant

Phylogenetic analysis was performed using Jalview (version 2.11.4.0) to assess the conservation of *C. elegans* fibulin-1 EGF repeat 5 relative to EGF repeat bearing proteins including the human fibulin-1 EGF repeat 5, factor IX EGF repeat 1 and fibrillin EGF repeat 12^59, 71^. The Jalview pairwise alignment tool was used to compare the amino acid sequences in these EGF repeats. *C. elegans* fibulin-1 EGF repeat 5 contains the six conserved cysteine residues that are characteristic of EGF repeats (Fig. 8a). To disrupt fibulin-1 association via EGF repeat 5, the cysteine (C) residue at amino acid position 413 (red) was substituted with a phenylalanine (F) residue via site-directed mutagenesis. Please refer to Supplementary Table 5 for oligonucleotide sequences.

### Construction of RNAi plasmids

RNAi constructs in this study were obtained from the Ahringer laboratory and Vidal laboratory RNAi libraries. RNAi constructs targeting *mNeonGreen* and *fem-3* were constructed in the T444T RNAi vector^72^. Please refer to Supplementary Table 5 for oligonucleotide sequences.

### RNAi experiment

Bacteria (HT115; DE3) harboring RNAi vectors were cultured in LB media that contains the selection marker 100 mg/mL ampicillin for 16 hours at 37°C. Double-stranded RNAi expression was induced by 1 mM Isopropyl ß-D-1-thiogalactopyranoside (IPTG) for one hour at 37°C. Bacteria cultures were plated on NGM plates that were treated with 1 mM IPTG and 100 mg/ml ampicillin. Plates were dried at 25°C for 8 hours before use.

To avoid spermathecal developmental defects, L1 worms were first grown on control plates (L4440 or T444T) to permit spermathecal development for 40 hours before they were washed and transferred onto feeding RNAi plates at the L4 stage (after spermathecal development is completed). Animals were subjected to feeding RNAi treatment for 24 hours (day 1 adult), 36 hours (day 1.5 adult), and 48 hours (day 2 adult). RNAi knockdown efficiencies (Supplementary Table 3) were determined via fluorescence microscopy in day 2 adult animals.

Germline-specific RNAi experiments were performed using the *C. elegans DCL569* strain^56^. This strain harbors a loss of function mutation in *rde-1*, an argonaute protein required for RNAi. Expression of *rde-1* specifically in the germline (*sun-1p::rde-1::sun-1* 3’ UTR) restores RNAi only in the germline^56^. L1 worms were grown on control plates for 40 hours before they were subjected to *fbl-1* feeding RNAi at the L4 stage. RNAi knockdown efficiencies (Supplementary Table 3) were determined via fluorescence microscopy in day 2 adult animals when analysis on ovulation and the spermatheca was performed. All experiments were carried out at 20°C.

Transgenerational *fem-3* RNAi was performed to inhibit ovulation. FEM-3 is required for sperm development in hermaphrodites, and in the absence of sperm, oocytes are not ovulated. Worms were grown on *fem-3* feeding RNAi plates for three generations for complete inhibition of sperm development, and then L4 worms were transferred onto either *fem-3* RNAi, *fbl-1* RNAi or combinatorial *fem-3* and *fbl-1* RNAi plates. The inhibition of ovulation was confirmed by assessing the absence of embryos on the *fem-3* RNAi plates. All experiments were performed at 20°C.

### Ovulation rate assessment

To assess ovulation rate, ten L4 worms were singled onto either control or *fbl-1* feeding RNAi plates. These worms were then transferred onto fresh plates at days 1, 1.5, 2, 2.5, 3.5 and 4.5 of adulthood. The number of ovulation events at each time point was determined by counting the number of progeny that were found on each plate two days after the parent worm had been transferred.

### Light microscopy

Confocal fluorescence microscopy experiments were performed with an upright Axio Imager.A1 (Carl Zeiss) that was mounted with a CSU-10 or W1 spinning disk confocal head (Yokogawa). Imaging experiments were performed either with a 40× Plan-Apochromat (NA 1.40) oil immersion objective or a 63× Plan-Apochromat (NA 1.40) (Carl Zeiss) oil immersion objective. Images were acquired using a scientific complementary metal-oxide semiconductor (CMOS) camera (ORCA-fusion; Hamamatsu) and a charged-coupled device camera (EM-CCD) (ImageEM; Hamamatsu). Images were acquired with a z-step size of 0.37 µm at room temperature.

To visualize the organization of fibulin-1 and type IV collagen in the spermathecal BM, confocal fluorescence microscopy experiments were performed on a Nikon Ti2-E inverted microscope equipped with a CSU-W1 spinning disk confocal head (Yokogawa). Images were acquired using a 100× SR HP Plan-Apochromat silicon objective (NA 1.35) and Hamamatsu Fusion BT CMOS camera. Photobleaching experiments were conducted with an iLas2 targeted laser system (BioVision) that was equipped with an Omicron Lux 60mW 405 nm continuous wave laser and controlled by the MetaMorph software. To assess BM stretching, we photobleached a spot (diameter: 5 µm) on the resting spermathecal BM (type IV collagen α2 chain; LET-2::mRuby) in day 1-2 adult animals. Photobleaching was limited to spermathecal BM surfaces that were devoid of folds. A timelapse imaging experiment was then conducted to track the photobleached spot through ovulation. Confocal fluorescence z-stacks images that span across the spermathecal BM surface (total z-stack range: 3.70 µm; z-step size: 0.37 µm) were acquired. The imaging frame interval was 10 seconds.

For single timepoint snapshots, worms were anaesthetized in 0.01 M sodium azide, mounted on 5% noble agar, and covered with a #1.5 cover slip for imaging. For imaging time course experiments, worms were anaesthetized in 5 mM lemavisole, mounted on 5% noble agar, and covered with a #1.5 cover slip for imaging. Cover slips were then sealed with VALAP to prevent sample evaporation. Each imaging time course experiment was limited to 1.5 hours.

### Image and statistical analyses

#### Quantification of BM/spermathecal volume, length and radius

The BM/spermathecal volume was quantified using a semiautomated strategy that utilized the FIJI (version 1.54f) macro scripting language and plugins^73^. Confocal z-stacks (pixel size: 0.16 µm; z-step size: 0.50 µm) of spermathecas (visualized with mCherry::moeABD) in control and *fbl-1* RNAi treated animals were acquired. To minimize the effects of out-of-focus blur, only spermathecas that were positioned near the cover slip side were imaged. The total fluorescence intensity of the spermatheca, which was centered within a 72 × 72 µm (451 × 451 pixel) box, was quantified. The background fluorescence intensity that was adjacent to the spermatheca was sampled and subtracted from the total fluorescence intensity. Images were then segmented and binarized via the Triangle thresholding method. The accuracy of image segmentation was manually inspected and corrected for each z-plane. The FIJI plugin Particle Analysis 3D was used to quantify the spermathecal volume. The accuracy of this semiautomated volume analysis strategy was validated by comparing all volume measurements against an alternate volume measurement method, which assumes that the spermatheca adopts an ellipsoid-like shape. The length and radius of each spermatheca were measured (Fig. 1e,f) and applied in the ellipsoid volume formula to approximate the volume of the spermatheca. The discrepancy in the spermathecal volume derived by both methods was confirmed to be under 10%. The length of each spermatheca (mCherry::moeABD) was measured (Fig. 1c) by manually tracing each spermatheca from the distal valve to the proximal valve. For radius measurement, it was sampled at the mid-section of the spermatheca.

#### Spermathecal BM stretching analysis

The surface area of the photobleached spot on resting and stretched spermathecal BMs was measured using the FIJI freehand selections tool (version 1.54f). The extent of BM stretching was calculated by dividing the surface area of the photobleached spot in the stretched state by the surface area of the photobleached spot in the resting state of the same spermathecal BM. The height and width of the photobleached spot were measured using the Line and Measure tools. The aspect ratio of the photobleached spot was quantified by dividing the spot width by its height (Extended Data Fig. 1b). All surface area measurements were limited to photobleached spots that were positioned on BM surfaces that were oriented parallel to the imaging plane and were devoid of BM folds (Fig. 1g).

#### Analyses of BM composition and RNAi knockdown efficiency

The relative level of each mNG-tagged BM component at the spermathecal BM was quantified using a semiautomated strategy that utilizes FIJI (version 1.54f). Using confocal z-stacks (pixel sizes: 0.16 µm or 0.40 µm; z-step size: 0.37 µm), the fluorescence intensity of each mNG-tagged BM component was quantified from the spermatheca that was positioned closer to the cover slip side. The total fluorescence intensity within a 1.1 × 1.1 µm box at the spermathecal BM was measured. The total background fluorescence intensity in the extracellular fluid surrounding the spermathecal BM was determined using a box of the same size. The net total fluorescence intensity signal was calculated by subtracting the local background fluorescence intensity signal from the total fluorescence intensity signal at the spermathecal BM. Two regions along the spermathecal BM were sampled. Measurements were taken from regions across the spermathecal BM surfaces that were devoid of spermathecal folds and were not in contact with BMs of adjacent tissues. To account for variability between imaging experiments, the average fluorescence intensity of each mNG-tagged BM components was normalized against the average fluorescence intensity of laminin (LAM-2::mNG), which served as a control that was included in all imaging experiments. The same image analysis strategy was applied to determine the abundance of BM proteins in the proximal gonadal BM and uterine BM and assess the knockdown efficiency of tagged BM proteins.

#### Analysis of oocyte fibulin-1 levels during ovulation

Fluorescence timelapse images of ovulation in day 1-2 adult animals expressing mNG::FBL-1 were acquired. Oocyte fibulin-1 levels were quantified using FIJI (version 1.54f). An outline of the oocyte was manually traced, and the total fluorescence intensity was quantified using the Measure tool. Background fluorescence intensity was quantified from outside the worm (camera background fluorescence intensity) using the same oocyte outline. The net fluorescence intensity was determined by subtracting the background fluorescence intensity from the total fluorescence intensity. The same measurement strategy was applied across all time points.

#### Analysis of zygote dimensions

The dimensions of control and *fbl-1* RNAi treated zygotes were measured using FIJI (version 1.54f). Confocal z-stacks images of animals expressing fluorescently tagged type IV collagen α2 chain (LET-2::mRuby) were acquired. As the uterine BM clearly marks the outline of zygotes, zygote dimensions were approximated using the uterine BM. Analysis was limited to the zygote that was positioned adjacent to the spermatheca. Length and width measurements were acquired from the image z-plane that bisects the midsection of the zygote.

#### FRAP analysis

FRAP analyses on fibulin-1 (mNG::FBL-1) and type IV collagen α1 chain (EMB-9::mRuby) were performed on day 1 and 2 adult animals. FRAP was conducted on regions of the spermathecal BM that did not overlap with adjacent BMs. Prior to photobleaching, the total fluorescence intensity within a specific region of the spermathecal BM was measured using a 2.4 × 0.8 µm (15 × 5 pixels) box. The background fluorescence intensity was sampled from the extracellular fluid adjacent to the spermathecal BM. The net fluorescence intensity of the spermathecal BM was calculated by subtracting the background fluorescence intensity from the total fluorescence intensity at the spermathecal BM. Upon photobleaching, the recovery in fluorescence intensity was assessed using the same fluorescence intensity quantification method. To determine the extent of photobleaching from timelapse imaging, the total fluorescence intensity at the worm hypodermal BM was measured using the same size box. Background fluorescence intensity was sampled from the worm exterior that was adjacent to the hypodermal BM. The net fluorescence intensity at the hypodermal BM was calculated by subtracting the background fluorescence intensity from the total fluorescence intensity at the hypodermal BM. The fraction of fluorescence intensity recovery was quantified by normalizing the net fluorescence intensity during the recovery phase against the net fluorescence intensity of the protein of interest prior to photobleaching. The fluorescence intensities of mNG::FBL-1 and mRuby::EMB-9 at the spermathecal BM were tracked over a duration of 10 min.

#### Actin organization analysis

F-actin organization was assessed by measuring the distance between individual actin bundles (mCherry::moeABD ) using FIJI (version 1.54f). For each pair of actin bundles, a perpendicular line that bisects them was manually drawn and the intervening distance was measured. Ten regions across the spermathecal actin were sampled per animal. Due to the irregular shape of the spermathecal surface, analysis was limited to spermathecal actin bundles that were oriented parallel to the imaging plane.

#### Fibulin-1 and spermathecal myoepithelial cells colocalization analysis

To determine if fibulin-1 is present within the spermathecal myoepithelial cells, confocal z-stack images of day 2 adult animals expressing fibulin-1 (mNG::FBL-1) and spermathecal actin (mCherry::moeABD) were acquired. Next, the image plane that bisects the mid-section of the spermatheca was extracted for processing. Each fluorescence channel was subjected to local background subtraction and then normalized to the brightest fluorescence intensity value. The fluorescence intensity distributions of mNG::FBL-1 and mCherry::moeABD that span across the spermathecal BM and tissue were obtained using the FIJI Plot Profile plugin. To compare the distribution of fibulin-1 and actin between resting and stretched spermathecas, the fluorescence intensity distributions that were acquired from each spermatheca were aligned based on the brightest actin fluorescence intensity.

#### Ovulation duration analysis

Ovulation duration was determined using timelapse movies of control and *fbl-1* RNAi treated animals (day 1 and 2 adults). Each ovulation window is defined as the time between when an oocyte first enters the spermatheca until the time when the zygote exits. The duration of ovulation was quantified by multiplying the number of imaging frames during the ovulation window with the imaging frame interval of 1 minute.

#### Fibulin-1 and type IV collagen organization and colocalization analyses

Confocal z-stacks of fibulin-1 (mNG::FBL-1) and type IV collagen α2 chain (LET-2::mRuby) within the BM of the spermatheca were acquired. All images were subjected to an image preprocessing protocol that included image denoising using the Nikon NIS-Elements Denoise.ai software. From this preprocessing, a single z-plane that spans across the spermathecal BM surface was extracted from the image z-stack. Only flat BM surfaces were used for analysis. The organization of fibulin-1 and type IV collagen in both resting and stretched BMs was assessed by measuring the distance between fibril/puncta using the Line tool in FIJI. To account for heterogeneity of the network, ten random regions across the BM surface of 6.7 x 6.7 µm box was sampled per animal. For each pair of fibrils/puncta, a perpendicular line that bisects them was manually drawn and the intervening distance was measured.

To assess the colocalization between the fibulin-1 and type IV collagen networks, fluorescence images were first preprocessed by local background subtraction, normalized to the brightest fluorescence intensity value, and segmented to generate a skeletonized image. Using the skeletonized image as a landmark, the fluorescence distribution of each protein was determined using the FIJI Plot Profile plugin and plotted for comparison (Extended Data Fig. 7p,q). Co-occurrence analysis was conducted to quantify the proportion of fibril/puncta that contained fibulin-1, type IV collagen, or both proteins in resting and stretched spermathecal BMs in day 2 adult animals.

#### Assessment of gaps in the type IV collagen network

To quantify the size of gaps present in the type IV collagen network across the spermathecal BM surface, the FIJI Line tool was used to measure the width of the gap in type IV collagen α2 chain (LET-2::mRuby) fluorescence signals. The width of each gap was sampled twice and averaged. To ensure consistency of measurements, analysis was only performed on regions of spermathecal BM surfaces that were oriented parallel to the imaging plane and that they were positioned away from the edge of the spermathecal BM. The assessment of type IV collagen gaps in BMs that were induced to stretched via *fln-1* RNAi treatment^54^ were limited to spermathecas that retained two oocytes.

#### Statistical analysis

All datasets were assessed for normal distribution using Shapiro-Wilk normality test. Student’s *t*-test was performed on normally distributed datasets. *Mann-Whitney* test was performed on

datasets that do not conform to normal distribution. Tests for significance were two-tailed and unpaired unless otherwise specified. All error bars indicate SD. Statistical significance was set at p value < 0.01. All analyses were performed on samples obtained from three independent experiments.

## Extended Data Figure legends

**Extended Data Figure 1.**
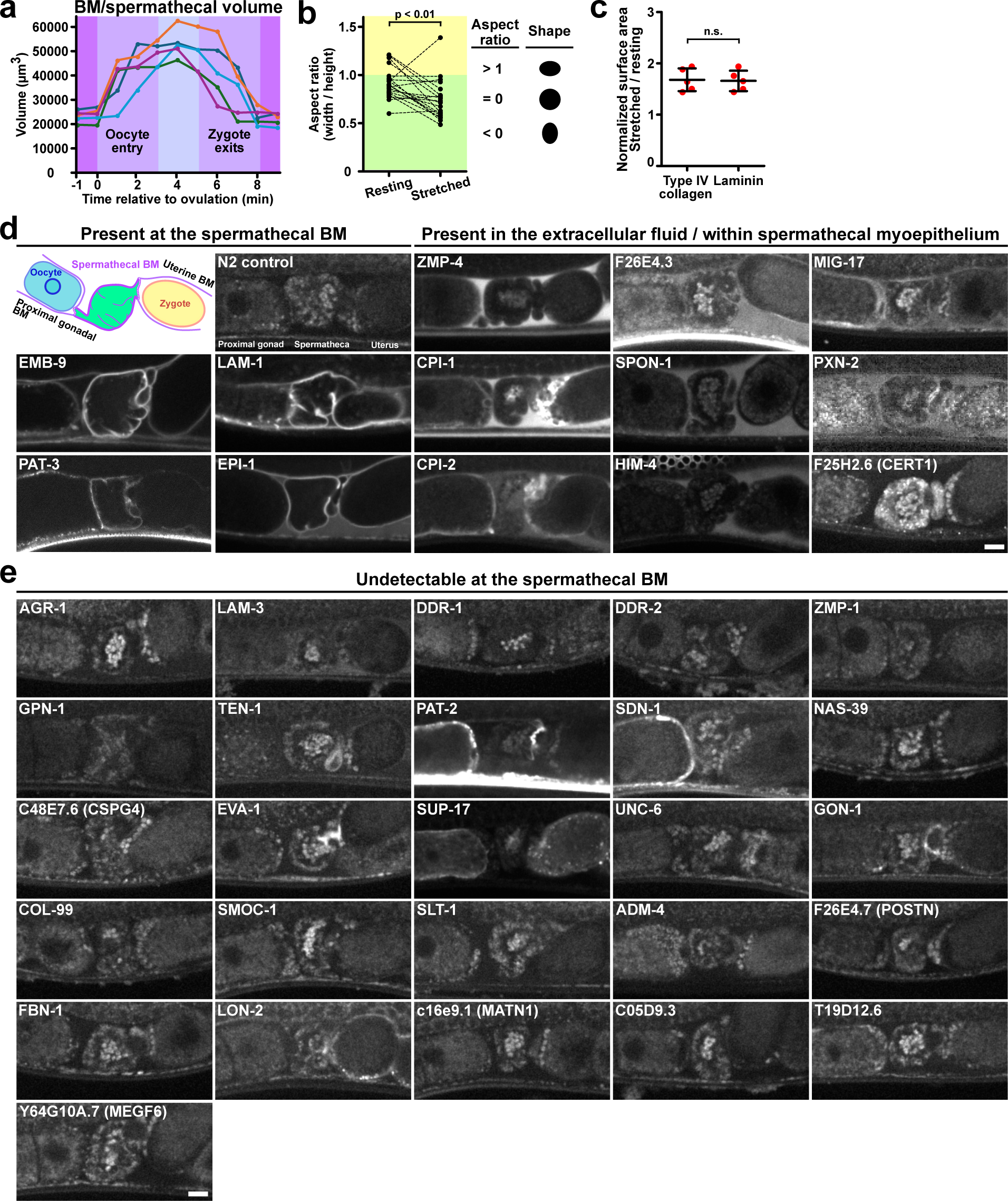
Characterization of the BM/spermathecal volume, stretching, and composition. **a**, Quantification of the BM/spermathecal volume through the course of an ovulation event. Each line represents the BM/spermathecal volume of an animal (n = 5 animals). **b,** (*Left*) Quantification of the aspect ratio (width/height) of each photobleached spot. (*Right*) A schematic diagram showing the representative shapes of photobleached spots (n = 20 animals). **c,** Quantification of the normalized surface area of photobleached spots in resting and stretched BMs that were visualized either by type IV collagen α2 chain (LET-2::mRuby) or laminin γ1 chain (LAM-2::mRuby; n = 5 animals; *Mann-Whitney* test; mean ± SD). **d,** (*Top left*) A schematic diagram showing the continuous BM that enwraps the proximal gonad, spermatheca and uterus. Fluorescence images (single z-slice) showing the localizations of endogenously mNG tagged BM proteins in the spermathecal BM, extracellular fluid, or within the spermathecal myoepithelium (intracellular). All fluorescence images were uniformly contrasted and compared against wild-type day 1 adult animals (N2) that do not express mNG (n = 5 animals). **e,** Fluorescence images (single z-slice) showing endogenously mNG tagged BM proteins that were not detected in the spermathecal BM (n = 5 animals). All data is representative of 3 independent biological repeats. Scale bars, 10 µm.

**Extended Data Figure 2.**
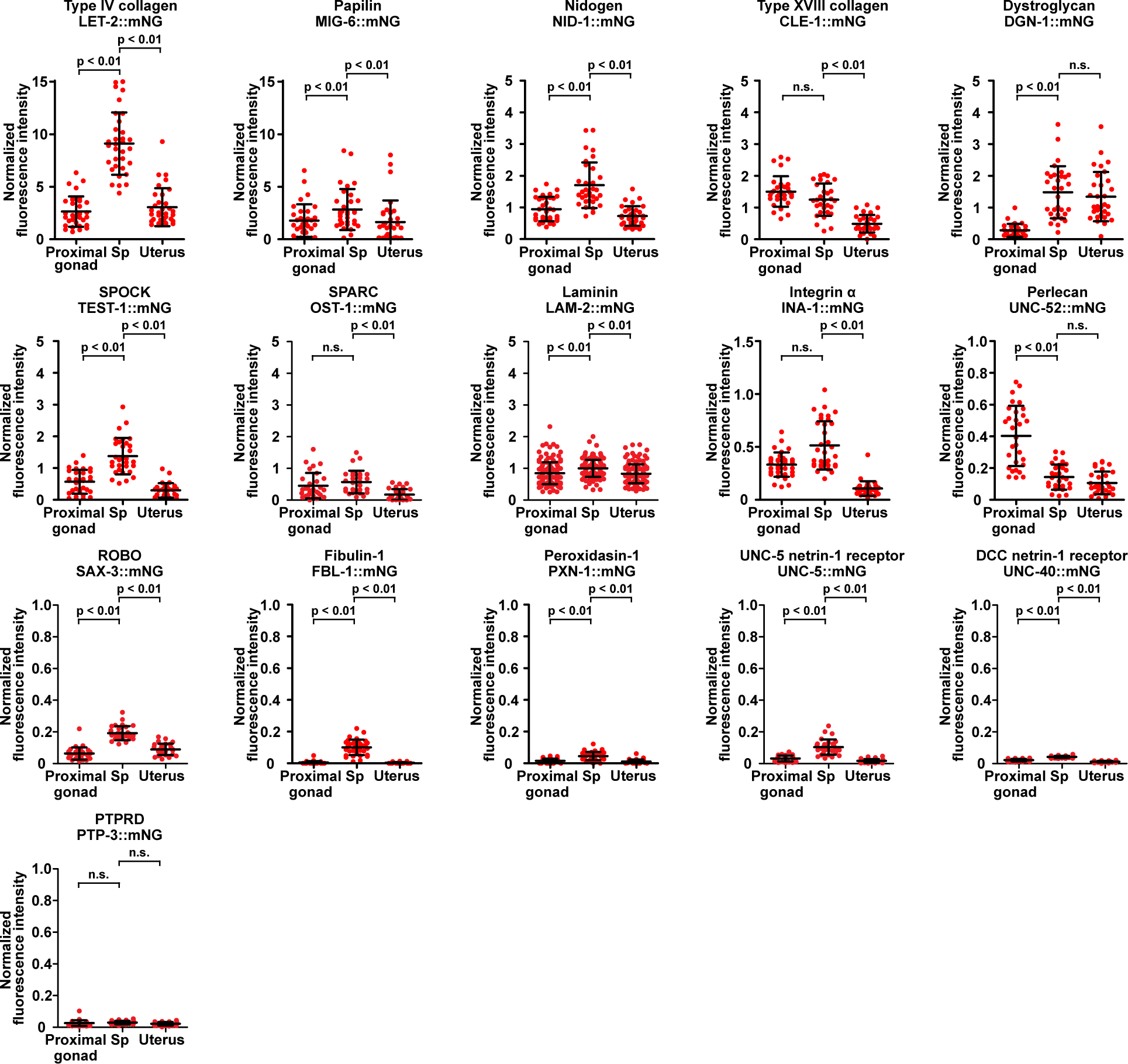
The spermathecal BM possesses a distinct composition. Quantification of the spermathecal BM protein levels during the resting state. The level of each BM protein in the proximal gonadal BM, spermathecal BM (Sp), and uterine BM was measured. All fluorescence intensities were normalized to the fluorescence intensity of LAM-2::mNG (laminin γ1 chain) at the spermathecal BM (n = 30-50 animals for all BM proteins, except for LAM-2::mNG where n = 145 animals; *Mann-Whitney* test; mean ± SD). All data is representative of 3 independent biological repeats.

**Extended Data Figure 3.**
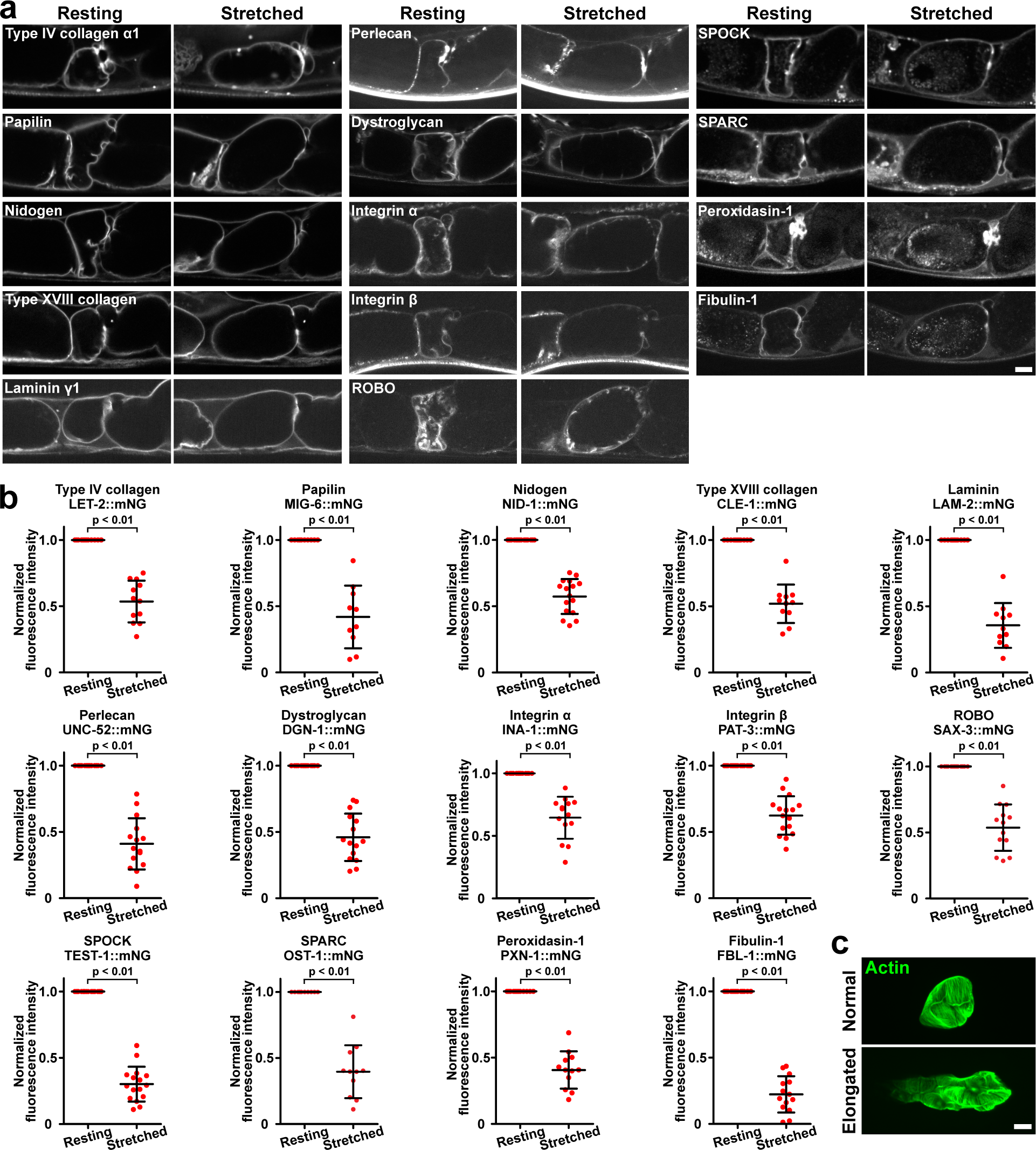
Spermathecal BM composition in resting and stretched BMs. **a**, Fluorescence images (single z-slice) showing endogenously mNG tagged BM proteins in resting and stretched spermathecal BMs. The fluorescence intensity of each BM protein was tracked through an ovulation event. All BM proteins represented in Fig. 2c were assessed, except for UNC-5::mNG, UNC-40::mNG and PTPRD::mNG as they were prone to photobleaching during live imaging (n = 10-16 animals). **b,** Quantification of spermathecal BM protein levels in resting and stretched BMs (n = 10-16 animals; Student’s *t*-test; mean ± SD). **c,** Maximum intensity z-projected fluorescence images showing a normal and an elongated spermatheca (actin, mCherry::moeABD, green). All data is representative of 3 independent biological repeats.

**Extended Data Figure 4.**
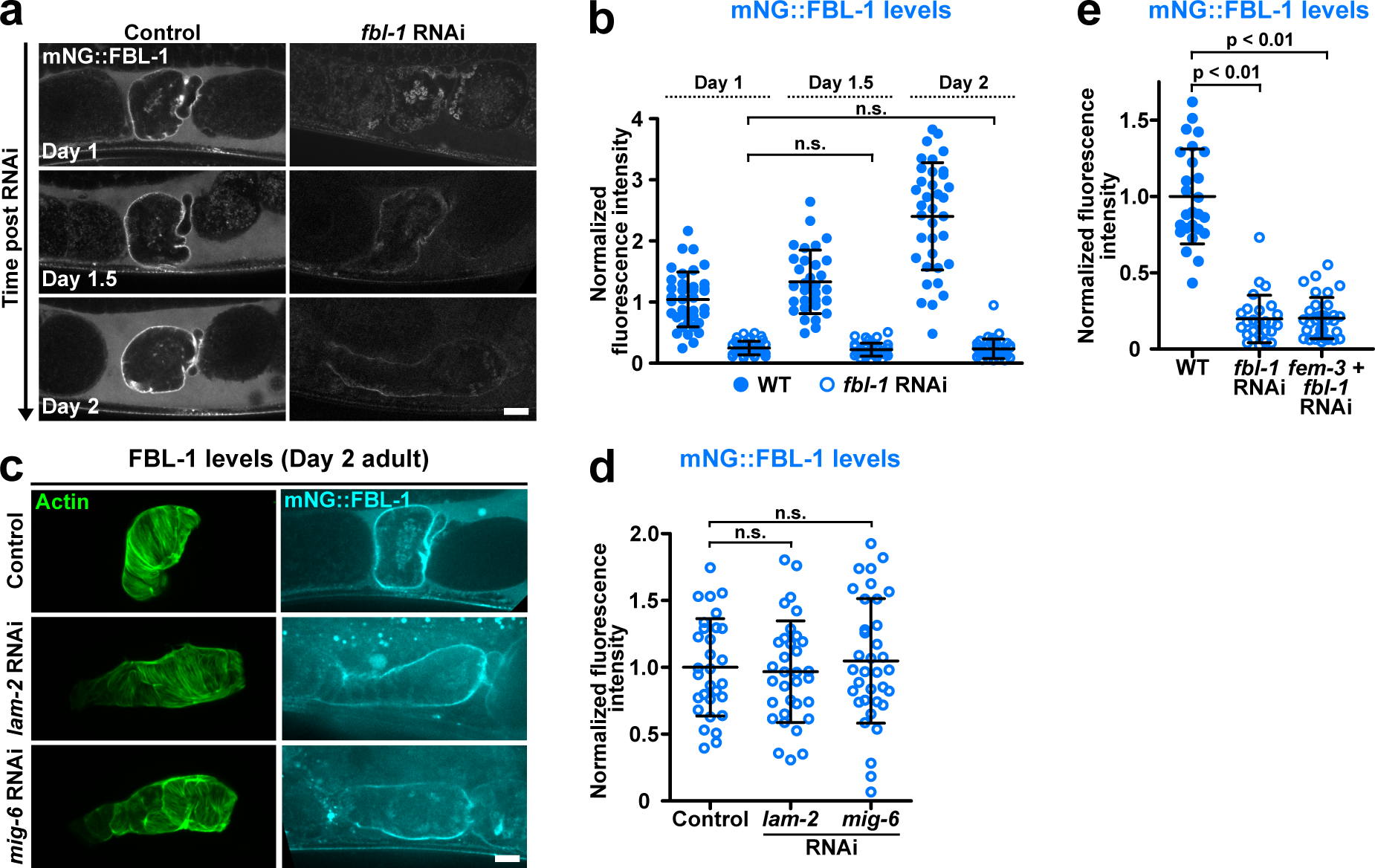
Fibulin-1 levels at the spermathecal BM during RNAi knockdown of laminin, papilin and fibulin-1 and inhibition of ovulation. **a**, Fluorescence images (single z-slice) of fibulin-1 (mNG::FBL-1, cyan) in control and *fbl-1* RNAi treated day 1-2 adult animals. **b,** Quantification of *fbl-1* RNAi knockdown efficiency in control and *fbl-1* RNAi treated day 1-2 adult animals (n = 31-41 animals; *Mann-Whitney* test; mean ± SD). **c,** Fluorescence images (single z-slice) showing the shape of the spermatheca (actin, mCherry::moeABD, green) and fibulin-1 (mNG::FBL-1, cyan) levels at the spermathecal BM in control, *lam-2* RNAi and papilin *(mig-6)* RNAi treated day 2 adult animals. **d,** Quantification of fibulin-1 levels in control, *lam-2* RNAi and papilin *(mig-6)* RNAi treated day 2 adult animals (Control, n = 30 animals; *lam-2* RNAi, n = 33 animals; *mig-6* RNAi, n = 35 animals; *Mann-Whitney* test; mean ± SD). **e,** Quantification of fibulin-1 (mNG::FBL-1) levels in control, *fbl-1* RNAi, and combinatorial *fem-3* and *fbl-1* RNAi treated day 2 adult animals (control, n = 26 animals; *fbl-1* RNAi, n = 26 animals; *fem-3* + *fbl-1* RNAi, n = 33 animals; *Mann-Whitney* test; mean ± SD). All data is representative of 3 independent biological repeats. Scale bars, 10 µm.

**Extended Data Figure 5.**
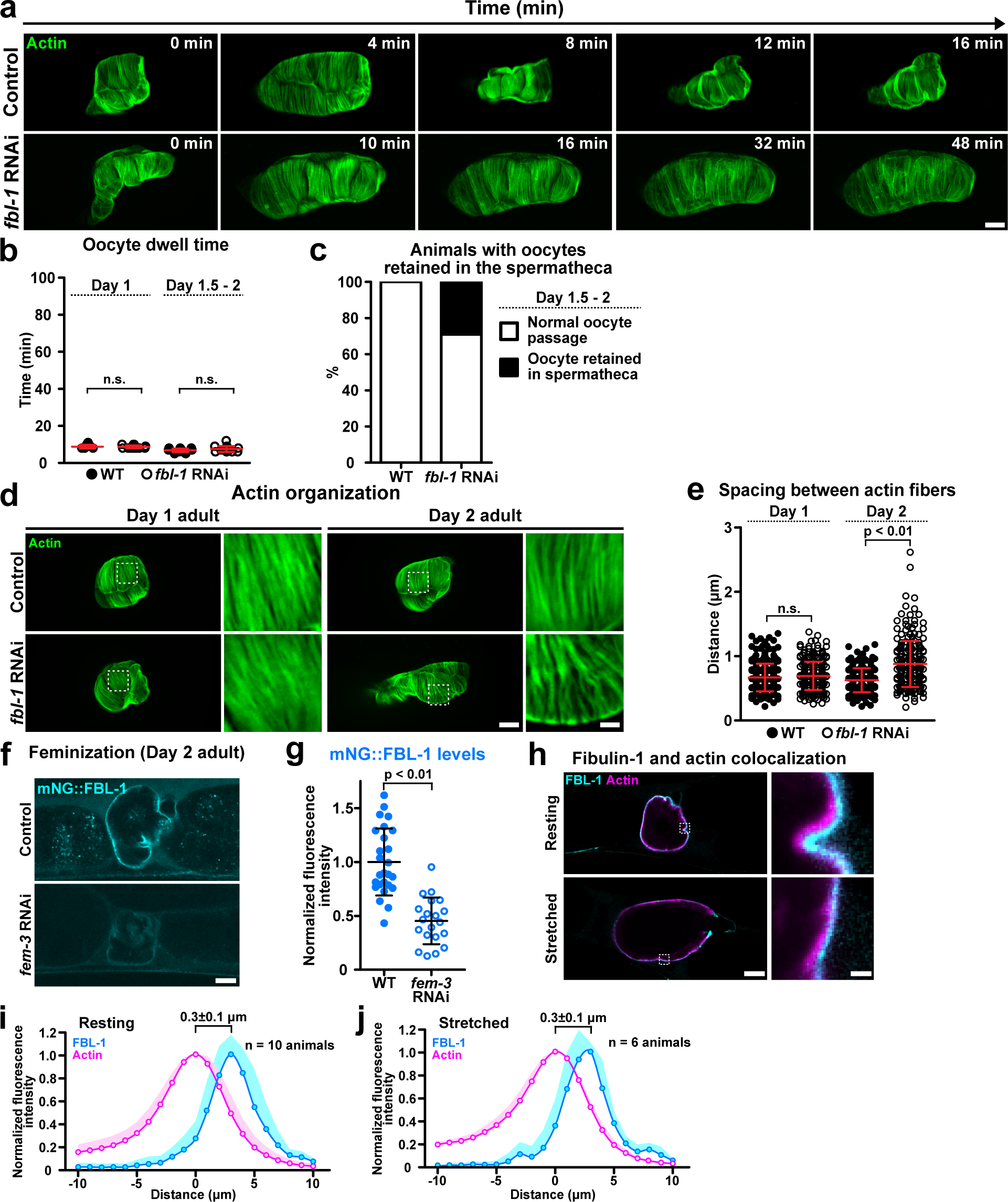
Fibulin-1 maintains spermathecal actin organization and spermathecal function. **a**, Maximum intensity z-projected fluorescence timelapse images of the spermatheca (actin, mCherry::moeABD, green) in control and *fbl-1* RNAi treated day 1.5 adult animals through an ovulation event. **b,** Quantification of oocyte dwell time in the spermatheca in control and *fbl-1* RNAi treated day 1-2 adult animals during ovulation (n = 10-16 animals; *Mann-Whitney* test; mean ± SD). **c,** Proportion of animals with oocytes that were retained in the spermatheca beyond the experimental time window of 80 min in control and *fbl-1* RNAi treated day 1.5-2 adult animals (control, n = 28 animals; *fbl-1* RNAi, n = 21 animals). **d,** Maximum intensity z-projected fluorescence images of the spermathecal actin (mCherry::moeABD , magenta) in control and *fbl-1* RNAi treated day 1 and 2 adult animals. **e,** Quantification of distance between actin bundles in control and *fbl-1* RNAi treated day 1 and 2 adult animals (n = 20-29 animals; *Mann-Whitney* test; mean ± SD). **f,** Fluorescence images (single z-slice) of fibulin-1 (mNG::FBL-1, cyan) in control and *fem-3* RNAi treated day 2 adult animals. **g,** Quantification of fibulin-1 (mNG::FBL-1) levels in control and *fem-3* RNAi treated day 2 adult animals (control, n = 26 animals; *fem-3* RNAi, n = 20 animals; *Mann-Whitney* test; mean ± SD). **h,** Fluorescence images (single z-slice) showing the localization of fibulin-1 (mNG::FBL-1, cyan) relative to the spermatheca (actin, mCherry::moeABD , magenta). **i-j,** Quantification of fluorescence intensity distributions of fibulin-1 (cyan) and the spermathecal actin (mCherry::moeABD , magenta) in (**i**) resting and (**j**) stretched BMs (resting state, n = 10 animals; stretched state, n = 6 animals; mean + SD). All data is representative of 3 independent biological repeats. Scale bars, 1 (insets), 10 µm.

**Extended Data Figure 6.**
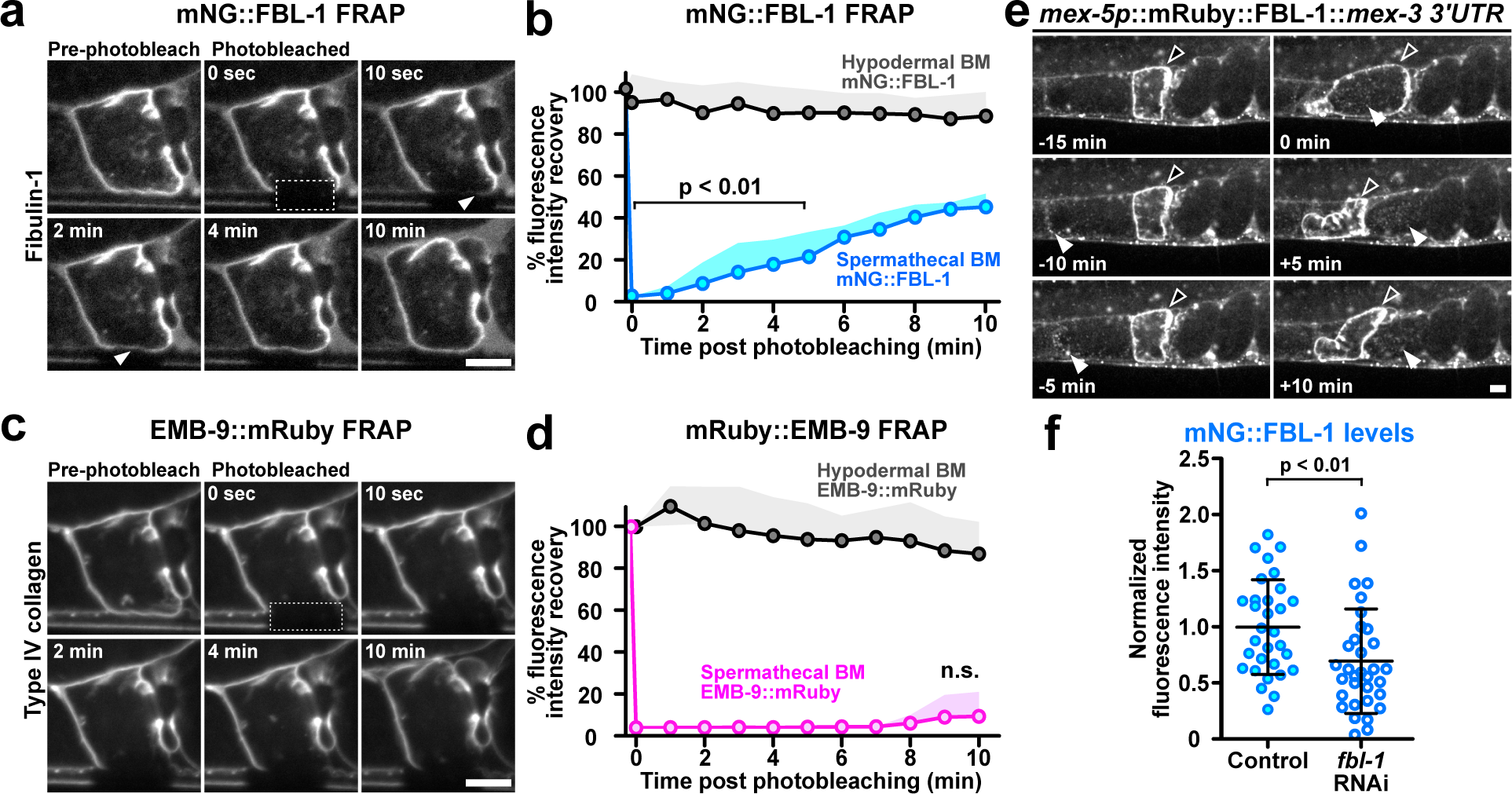
Fibulin-1 is secreted by ovulating oocytes and has rapid on-off association with the spermathecal BM. **a**, Fluorescence timelapse images (single z-slice) showing fibulin-1 (mNG::FBL-1, gray) FRAP (dotted rectangle, arrowhead) over a 10 min period. **b,** Quantification of mNG::FBL-1 FRAP (n = 5 animals; *Mann-Whitney* test; mean + SD). **c,** Fluorescence timelapse images (single z-slice) showing type IV collagen α1 chain (mRuby::EMB-9, gray) FRAP (dotted rectangle) over a 10 min period. **d,** Quantification of type IV collagen α1 chain (mRuby::EMB-9) FRAP (n = 5 animals; *Mann-Whitney* test; mean + SD). **e,** Fluorescence timelapse images (single z-slice) showing that oocyte expressed (*mex-5p::mRuby::fbl-1::mex-3 3’UTR*) fibulin-1 (mRuby::FBL-1, gray) is present in the spermathecal BM (outlined arrowhead) and within the oocyte/zygote (filled arrowhead). **f,** Quantification of fibulin-1 (mNG::FBL-1) levels in control and germline *fbl-1* RNAi treated day 2 adult animals (control, n = 31 animals; RNAi, n = 33 animals; *Mann-Whitney* test; mean ± SD). All data is representative of 3 independent biological repeats.

**Extended Data Figure 7.**
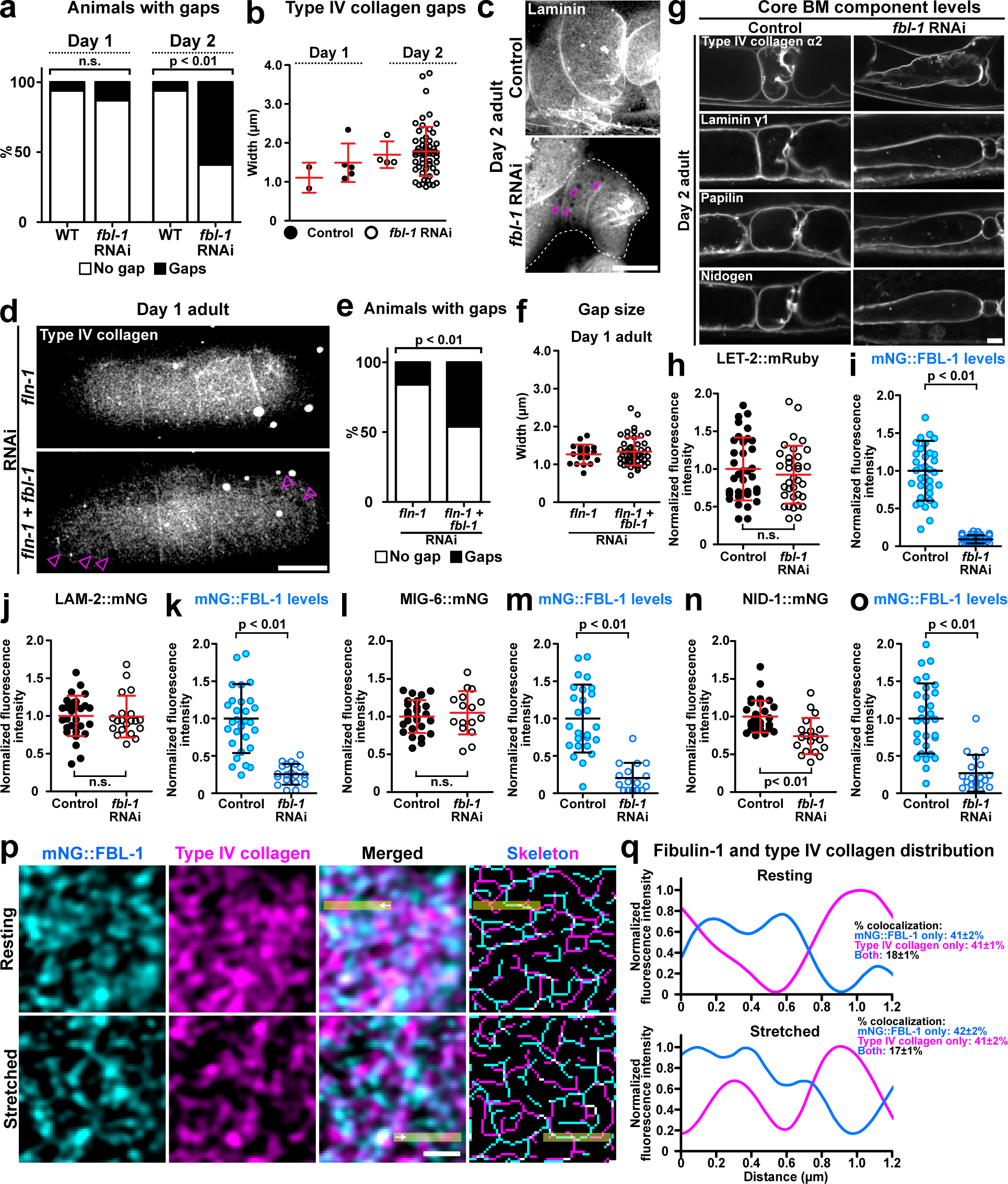
Fibulin-1 maintains the integrity, but not levels, of type IV collagen and laminin in the spermathecal BM. **a**, Proportion of animals with type IV collagen gaps in control and *fbl-1* RNAi treated animals (control day 1 adult, n = 24 animals; *fbl-1* RNAi day 1 adult, n = 22 animals; control day 2 adult, n = 25 animals; *fbl-1* RNAi day 2 adult, n = 27 animals; *Fisher’s exact* test). **b,** Quantification of type IV collagen gap widths in control and *fbl-1* RNAi treated animals (control day 1 adult, n = 24 animals; *fbl-1* RNAi day 1 adult, n = 22 animals, control day 2 adult, n = 25 animals; *fbl-1* RNAi day 2 adult, n = 27 animals; *Mann-Whitney* test; mean ± SD). **c,** Maximum intensity z-projected fluorescence images showing the surface view of the spermathecal BM (LAM-2::mNG, gray) in control and *fbl-1* RNAi treated day 2 adult animals where gaps were observed (magenta arrowheads). **d,** Maximum intensity z-projected fluorescence images showing the surface view of the spermathecal BM (LET-2::mRuby, gray) in filamin-1 (*fln-1*) and combinatorial *fln-1* + *fbl-1* RNAi treated day 1 adult animals where gaps were observed (magenta arrowheads). **e,** Proportion of animals with type IV collagen gaps (*fln-1* RNAi, n = 25 animals; *fln-1 + fbl-1* RNAi, n = 21 animals; *Fisher’s exact* test). **f,** Quantification of type IV collagen gap widths in *fln-1* and combinatorial *fln-1* + *fbl-1* RNAi treated day 1 adult animals (*fln-1* RNAi, n = 25 animals; *fln-1 + fbl-1* RNAi, n = 21 animals; *Mann-Whitney* test; mean ± SD). **g,** Fluorescence images (single z-slice) showing the core BM proteins type IV collagen α2 chain (LET-2::mRuby, gray), laminin γ1 chain (LAM-2::mNG, gray), papilin (MIG-6::mNG), and nidogen (NID-1::mNG) at the spermathecal BM in control and *fbl-1* RNAi treated day 2 adult animals. **(h-o)** Quantification of BM components after *fbl-1* RNAi knockdown (each data point represents the averaged fluorescence intensity sampled from each animal; *Mann-Whitney* test. Mean ± SD). **p,** Fluorescence (single z-slice) and skeletonized images showing the fibulin-1 (mNG::FBL-1, cyan) and type IV collagen α2 chain (LET-2::mRuby, magenta) networks. The fluorescence intensity distributions (yellow rectangle with arrows) were assessed. **q,** Fluorescence intensity distributions of fibulin-1 (mNG::FBL-1, cyan) and type IV collagen α2 chain (LET-2::mRuby, magenta). Co-occurrence analysis showing the proportions of fibril/puncta that contained fibulin-1, type IV collagen, or both proteins in resting and stretched spermathecal BMs (resting state n = 90 pairs of fibrils from 9 animals; stretched state n = 140 pairs of fibrils from 14 animals). All data is representative of 3 independent biological repeats. Scale bars, 10 µm (**c,d,g**) and 1 µm (**p**).

**Extended Data Figure 8.**
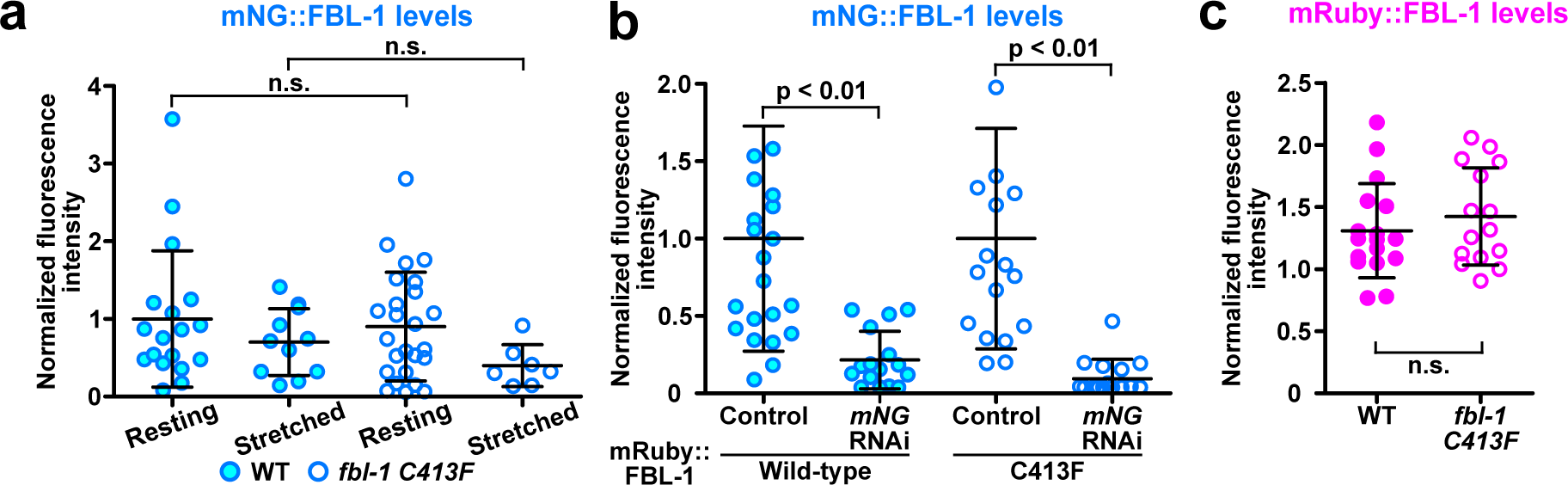
Exogenous wild-type and C413F mutant fibulin-1 levels. **a**, Quantification of mNG tagged wild-type and C413F mutant fibulin-1 (mNG::FBL-1) levels in resting and stretched spermathecal BMs in day 1-2 adult animals (wild-type resting state, n = 18 animals; wild-type stretched state, n = 11 animals; C413F mutant resting state, n = 25 animals; C413F mutant stretched state, n = 7 animals; *Mann-Whitney* test; mean ± SD). **b,** Quantification of endogenous mNG::FBL-1 levels (control + exogenous mRuby::FBL-1, n = 23 animals; *mNG* RNAi + exogenous mRuby::FBL-1, n = 16 animals; control + exogenous mRuby::FBL-1 C413F mutant, n = 18 animals; *mNG* RNAi + mRuby::FBL-1 C413F mutant, n = 16 animals; *Mann-Whitney* test. Mean ± SD). **c,** Quantification of exogenous wild-type and C413F mutant fibulin-1 (mRuby::FBL-1) levels (wild-type, n = 16 animals; C413F mutant, n = 15 animals; Student’s *t*-test; mean± SD). All data is representative of 3 independent biological repeats.

## Supplementary Tables

**Supplementary Table 1.**
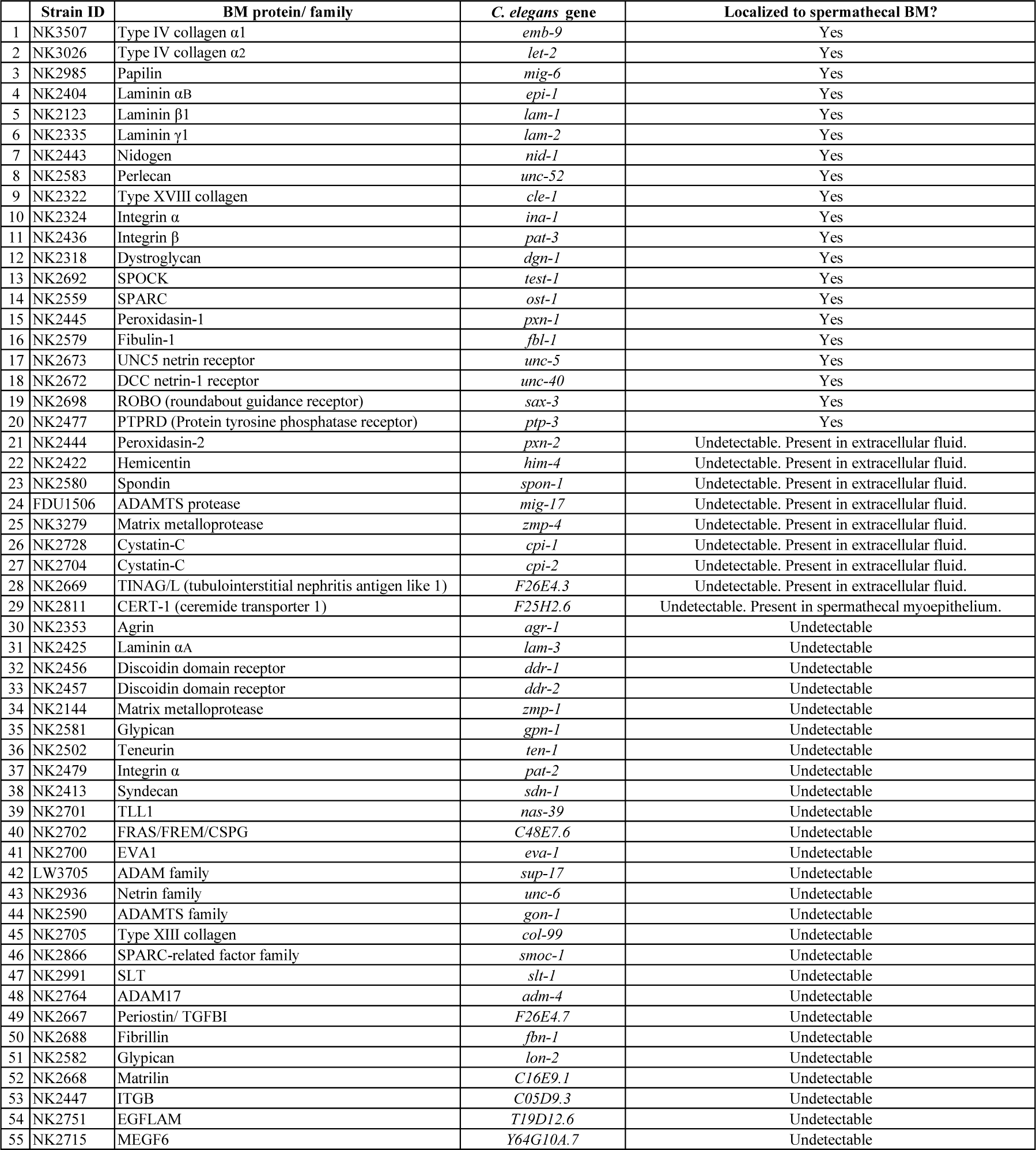
Spermathecal BM protein visual screen.

**Supplementary Table 2.** RNAi screen.

|  | Percentage of elongated spermatheca (%) |  |  |  |  |  |
| --- | --- | --- | --- | --- | --- | --- |
|  | BM protein | *C. elegans* genes | 24 h post RNAi |  | 48 h post RNAi |  |
|  |  |  | % | n | % | n |
| 1 | Control (empty RNAi vector) | Control vector (L4440/ T444T) | 0.0 | 90 | 10.0 | 90 |
| 2 | Type IV collagen $\alpha 1$ | *emb-9* | 96.7 | 30 | 100.0 | 30 |
| 3 | Papillin | *mig-6* | 93.3 | 30 | 100.0 | 30 |
| 4 | Laminin $\gamma 1$ | *lam-2* | 86.7 | 30 | 100.0 | 30 |
| 5 | Nidogen | *nid-1* | 46.7 | 30 | 63.3 | 30 |
| 6 | Perlecan | *unc-52* | 13.3 | 30 | 10.0 | 30 |
| 7 | Type XVIII collagen | *cle-1* | 3.3 | 30 | 13.3 | 30 |
| 8 | Integrin $\alpha$ | *ina-1* | 6.7 | 30 | 10.0 | 30 |
| 9 | Dystroglycan | *dgn-1* | 10.0 | 30 | 13.3 | 30 |
| 10 | SPOCK | *test-1* | 6.7 | 30 | 16.7 | 30 |
| 11 | SPARC | *ost-1* | 3.0 | 30 | 3.0 | 30 |
| 12 | Peroxidasin-1 | *pxn-1* | 6.7 | 30 | 10.0 | 30 |
| 13 | Peroxidasin-2 | *pxn-2* | 0.0 | 30 | 3.0 | 30 |
| 14 | Hemicentin | *him-4* | 0.0 | 30 | 0.0 | 30 |
| 15 | Spondin-1 | *spon-1* | 6.0 | 30 | 10.0 | 30 |
| 16 | ADAMTS protease | *mig-17* | 6.0 | 30 | 0.0 | 30 |
| 17 | Matrix metalloprotease | *zmp-4* | 0.0 | 30 | 3.0 | 30 |
| 18 | Cystatin-C | *cpi-1* | 0.0 | 30 | 0.0 | 30 |
| 19 | Cystatin-C | *cpi-2* | 0.0 | 30 | 3.0 | 30 |
| 20 | CERT-1 (ceramide transporter 1) | *F25H2.6* | 3.0 | 30 | 6.0 | 30 |
| 21 | TINAG/L (tubulointerstitial nephritis antigen like 1) | *F26E4.3* | 3.0 | 30 | 6.0 | 30 |
| 22 | UNC5 netrin receptor | *unc-5* | 3.0 | 30 | 3.0 | 30 |
| 23 | DCC netrin-1 receptor | *unc-40* | 10.0 | 30 | 10.0 | 30 |
| 24 | ROBO (roundabout guidance receptor) | *sax-3* | 0.0 | 30 | 0.0 | 30 |
| 25 | PTPRD (Protein tyrosine phosphatase receptor) | *ptp-3* | 3.0 | 30 | 3.0 | 30 |
| 26 | Fibulin-1 | *fbl-1* | 30.0 | 30 | 96.7 | 30 |

\*Spermathecal shape was assessed in day 1 adults (24h post RNAi) and day 2 adults (48h post RNAi). Please refer to Extended Data Fig. 3c for representative images of control and elongated spermathecas.

**Supplementary Table 3.**
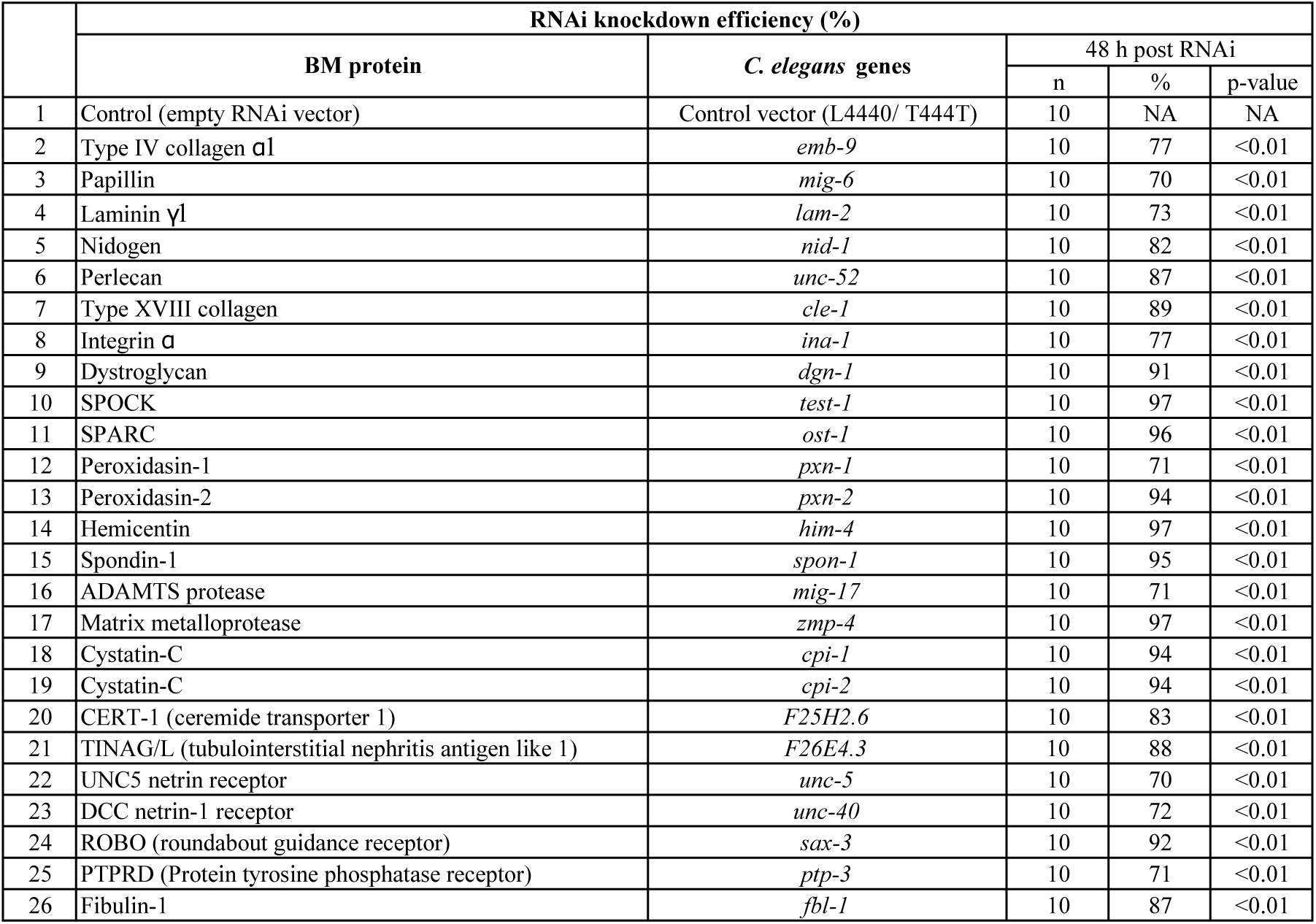
RNAi knockdown efficiency.

**Supplementary Table 4.**
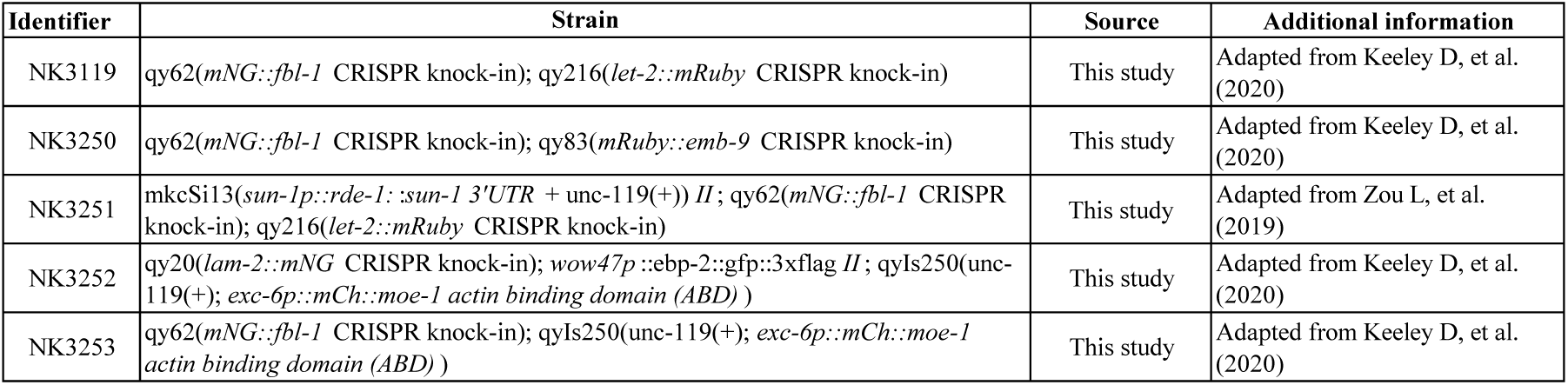
Strains.

**Supplementary Table 5.**
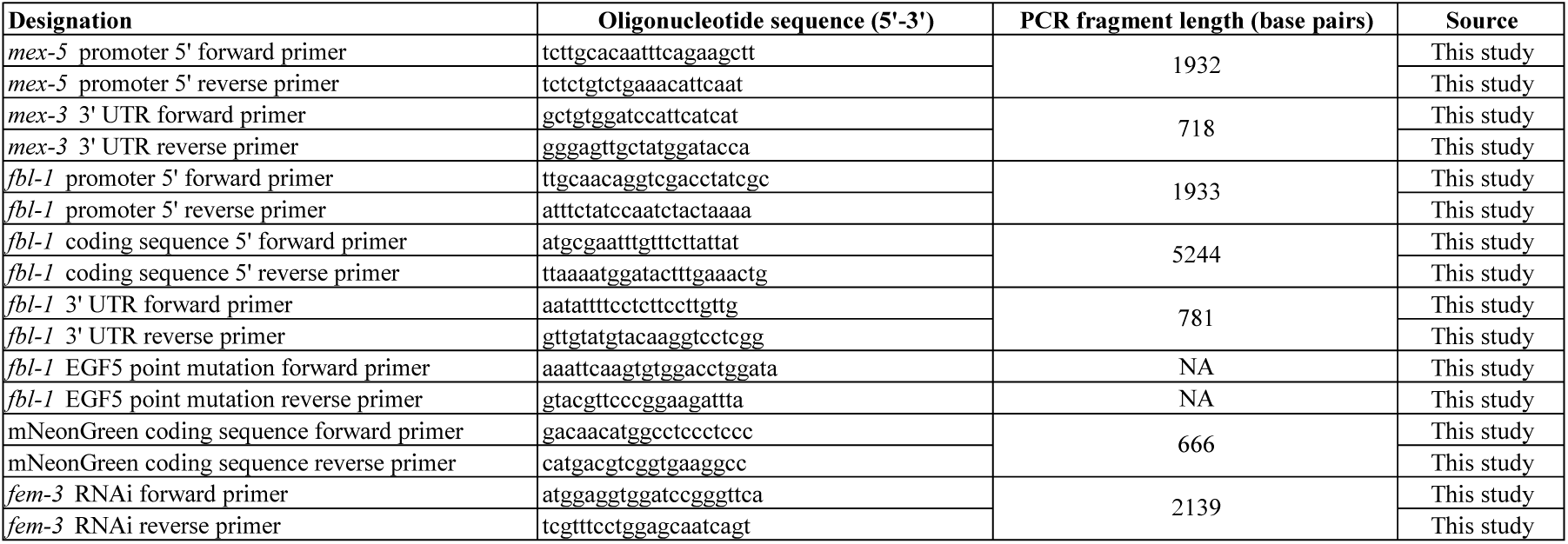
Oligonucleotide sequences.

## Supplementary Video legends

**Supplementary Video 1. The *C. elegans* spermatheca and its enveloping BM expand and recover during ovulation**. Fluorescence timelapse images showing a 3D projection of the spermathecal BM (laminin γ1 chain, LAM-2::mNG, magenta) and spermatheca (actin, mCherry::moeABD, green). Cells were visualized by a microtubule plus-end binding protein (EBP-3::GFP, magenta). Movie duration, 39 min. Imaging frame interval, 1.25 min. Framerate, 5 frames per second. n = 5 animals. Scale bar, 10 µm.

**Supplementary Video 2. Fibulin-1 maintains actin organization and spermathecal function.** Maximum intensity z-projected timelapse images showing spermathecal (actin, mCherry::moeABD, green) in control and *fbl*-1 RNAi treated day 1.5 adult animals. In the control animal, oocyte passage proceeded in a normal fashion over an eight-minute window. In *fbl-1* RNAi treated animal, the oocyte entered the spermatheca but failed to exit. In addition, actin bundles frayed and became spaced further apart. Movie duration, 16 min. Imaging frame interval, 1 min. Framerate, 4 frames per second. n = 5 animals. Scale bar, 10 µm.

**Supplementary Video 3. Ovulating oocytes are enriched with fibulin-1 as they transit the spermatheca**. Fluorescence timelapse images (single z-slice) showing endogenously expressed fibulin-1 (mNG::FBL-1; gray) in the ovulating oocytes, spermathecal BM and extracellular fluid over the course of two consecutive ovulation events. Oocyte fibulin-1 levels peaked as the oocyte entered the spermatheca. A rapid reduction in oocyte fibulin-1 levels ensued as the oocyte passed through the spermatheca. Movie duration, 90 min. Imaging frame interval, 1 min. Framerate, 4 frames per second. n = 6 animals Scale bar, 10 µm.

**Supplementary Video 4. Fibulin-1 turns over more rapidly than type IV collagen at the spermathecal BM**. Fluorescence timelapse images (single z-slice) showing FRAP of fibulin-1 (mNG::FBL-1, cyan) and type IV collagen α1 chain (EMB-9::mRuby, magenta). Movie duration, 10.3 min. Imaging frame interval, 10 sec. Framerate, 4 frames per second. n = 5 animals. Scale bar, 10 µm.

**Supplementary Video 5. Oocyte-expressed fibulin-1 is enriched at the spermathecal BM.** Fluorescence timelapse images (single z-slice) showing oocyte expressed fibulin-1 (mRuby::FBL-1, gray) is enriched in the oocyte and spermathecal BM. Movie duration, 90 min. Imaging frame interval, 1 min. Framerate, 4 frames per second. n = 10 animals Scale bar, 10 µm.

## Acknowledgement

We thank the Sherwood laboratory members for helpful discussions on this manuscript. We are thankful to Drs. Veronica Farmer and Amy S. Gladfelter (Gladfelter laboratory, Duke University) for their generosity in sharing microscopy resources and image analysis expertise. We thank Drs. Conor Sugden and Rachel Lennon (Lennon laboratory, University of Manchester) for insightful discussions on fibulin-1 patient mutations. We also thank Dr. Kerry Bloom (Bloom laboratory, University of North Carolina Chapel Hill) for providing feedback on this manuscript. This work was supported by the Jane Coffin Childs Memorial Fund for Medical Research and Duke Regeneration Center Postdoctoral Research Award to Accelerate Career Independence to A.W.J. Soh, and R35GM118049, R21OD032430 and Wellcome Trust Discovery Award (Ref: 227417/Z/23/Z) to D.R. Sherwood.

## References

1. Trębacz, H. & Barzycka, A. Mechanical Properties and Functions of Elastin: An Overview. Biomolecules 13 (2023).

2. Schmelzer, C.E.H. & Duca, L. Elastic fibers: formation, function, and fate during aging and disease. Febs j 289, 3704–3730 (2022).

3. Muiznieks, L.D. & Keeley, F.W. Molecular assembly and mechanical properties of the extracellular matrix: A fibrous protein perspective. Biochim Biophys Acta 1832, 866–875 (2013).

4. Gosline, J. et al. Elastic proteins: biological roles and mechanical properties. Philos Trans R Soc Lond B Biol Sci 357, 121–132 (2002).

5. Ishizaki, Y., Wang, J., Kim, J., Matsumoto, T. & Maeda, E. Contributions of collagen and elastin to elastic behaviours of tendon fascicle. Acta Biomater 176, 334–343 (2024).

6. Tsamis, A., Krawiec, J.T. & Vorp, D.A. Elastin and collagen fibre microstructure of the human aorta in ageing and disease: a review. J R Soc Interface 10, 20121004 (2013).

7. Najjar, S.S., Scuteri, A. &Lakatta, E.G. Arterial Aging. Hypertension 46, 454–462 (2005).

8. Sugawara, J., Hayashi, K., Yokoi, T. & Tanaka, H. Age-associated elongation of the ascending aorta in adults. JACC Cardiovasc Imaging 1, 739–748 (2008).

9. D’Errico, A. et al. Changes in the alveolar connective tissue of the ageing lung. An immunohistochemical study. Virchows Arch A Pathol Anat Histopathol 415, 137–144 (1989).

10. Górski, P., Białas, A.J. & Piotrowski, W.J. Aging Lung: Molecular Drivers and Impact on Respiratory Diseases—A Narrative Clinical Review. Antioxidants 13, 1480 (2024).

11. Jayadev, R. & Sherwood, D.R. Basement membranes. Curr Biol 27, R207–r211 (2017).

12. Khalilgharibi, N. & Mao, Y. To form and function: on the role of basement membrane mechanics in tissue development, homeostasis and disease. Open Biol 11, 200360 (2021).

13. Adamo, C.S., Zuk, A.V. & Sengle, G. The fibrillin microfibril/elastic fibre network: A critical extracellular supramolecular scaffold to balance skin homoeostasis. Exp Dermatol 30, 25–37 (2021).

14. Tohgasaki, T., Nishizawa, S., Kondo, S., Ishiwatari, S. & Sakurai, T. Long Hanging Structure of Collagen VII Connects the Elastic Fibers and the Basement Membrane in Young Skin Tissue. J Histochem Cytochem 70, 751–757 (2022).

15. Tiedemann, K. et al. Microfibrils at basement membrane zones interact with perlecan via fibrillin-1. J Biol Chem 280, 11404–11412 (2005).

16. Godwin, A.R.F. et al. The role of fibrillin and microfibril binding proteins in elastin and elastic fibre assembly. Matrix Biol 84, 17–30 (2019).

17. Li, H., Zheng, Y., Han, Y.L., Cai, S. & Guo, M. Nonlinear elasticity of biological basement membrane revealed by rapid inflation and deflation. Proc Natl Acad Sci U S A 118 (2021).

18. Salimbeigi, G., Vrana, N.E., Ghaemmaghami, A.M., Huri, P.Y. & McGuinness, G.B. Basement membrane properties and their recapitulation in organ-on-chip applications. Mater Today Bio 15, 100301 (2022).

19. Chang, J. & Chaudhuri, O. Beyond proteases: Basement membrane mechanics and cancer invasion. J Cell Biol 218, 2456–2469 (2019).

20. Yurchenco, P.D. & Ruben, G.C. Basement membrane structure in situ: evidence for lateral associations in the type IV collagen network. J Cell Biol 105, 2559–2568 (1987).

21. Yurchenco, P.D. & Ruben, G.C. Type IV collagen lateral associations in the EHS tumor matrix. Comparison with amniotic and in vitro networks. Am J Pathol 132, 278–291 (1988).

22. Borchiellini, C., Coulon, J. & Le Parco, Y. The function of type IV collagen during Drosophila muscle development. Mech Dev 58, 179–191 (1996).

23. Gupta, M.C., Graham, P.L. & Kramer, J.M. Characterization of alpha1(IV) collagen mutations in Caenorhabditis elegans and the effects of alpha1 and alpha2(IV) mutations on type IV collagen distribution. J Cell Biol 137, 1185–1196 (1997).

24. Pöschl, E. et al. Collagen IV is essential for basement membrane stability but dispensable for initiation of its assembly during early development. Development 131, 1619–1628 (2004).

25. Fidler, A.L., Boudko, S.P., Rokas, A. & Hudson, B.G. The triple helix of collagens - an ancient protein structure that enabled animal multicellularity and tissue evolution. J Cell Sci 131 (2018).

26. Jayadev, R. et al. A basement membrane discovery pipeline uncovers network complexity, new regulators, and human disease associations. bioRxiv, 2021.2010.2025.465762 (2021).

27. Ambade, A.S., Hassoun, P.M. & Damico, R.L. Basement Membrane Extracellular Matrix Proteins in Pulmonary Vascular and Right Ventricular Remodeling in Pulmonary Hypertension. Am J Respir Cell Mol Biol 65, 245–258 (2021).

28. Boland, E., Quondamatteo, F. & Van Agtmael, T. The role of basement membranes in cardiac biology and disease. Biosci Rep 41 (2021).

29. Dekkers, B.G.J., Saad, S.I., van Spelde, L.J. & Burgess, J.K. Basement membranes in obstructive pulmonary diseases. Matrix Biol Plus 12, 100092 (2021).

30. Nyström, A., Bornert, O. & Kühl, T. Cell therapy for basement membrane-linked diseases. Matrix Biol 57-58, 124-139 (2017).

31. Mahajan, D. et al. Role of Fibulins in Embryonic Stage Development and Their Involvement in Various Diseases. Biomolecules 11 (2021).

32. Kostka, G. et al. Perinatal lethality and endothelial cell abnormalities in several vessel compartments of fibulin-1-deficient mice. Mol Cell Biol 21, 7025–7034 (2001).

33. Argraves, W.S., Greene, L.M., Cooley, M.A. & Gallagher, W.M. Fibulins: physiological and disease perspectives. EMBO Rep 4, 1127–1131 (2003).

34. Tran, H., VanDusen, W.J. & Argraves, W.S. The self-association and fibronectin-binding sites of fibulin-1 map to calcium-binding epidermal growth factor-like domains. J Biol Chem 272, 22600–22606 (1997).

35. Sasaki, T. et al. Structural characterization of two variants of fibulin-1 that differ in nidogen affinity. J Mol Biol 245, 241–250 (1995).

36. Olin, A.I. et al. The proteoglycans aggrecan and Versican form networks with fibulin-2 through their lectin domain binding. J Biol Chem 276, 1253–1261 (2001).

37. McCarter, J., Bartlett, B., Dang, T. & Schedl, T. On the control of oocyte meiotic maturation and ovulation in Caenorhabditis elegans. Dev Biol 205, 111–128 (1999).

38. Kelley, C.A. & Cram, E.J. Regulation of Actin Dynamics in the C. elegans Somatic Gonad. J Dev Biol 7 (2019).

39. Ward, S. & Carrel, J.S. Fertilization and sperm competition in the nematode Caenorhabditis elegans. Dev Biol 73, 304–321 (1979).

40. Hall, R.L.a.D.H., Vol. 2024 (2009).

41. Kocsisova, Z., Kornfeld, K. & Schedl, T. Rapid population-wide declines in stem cell number and activity during reproductive aging in C. elegans. Development 146 (2019).

42. Keeley, D.P. et al. Comprehensive Endogenous Tagging of Basement Membrane Components Reveals Dynamic Movement within the Matrix Scaffolding. Dev Cell 54, 60–74.e67 (2020).

43. Schindler, A.J. & Sherwood, D.R. The transcription factor HLH-2/E/Daughterless regulates anchor cell invasion across basement membrane in C. elegans. Developmental Biology 357, 380–391 (2011).

44. De Las Heras, J.M., et al. The Drosophila Hox gene Ultrabithorax controls appendage shape by regulating extracellular matrix dynamics. Development 145 (2018).

45. Pastor-Pareja, J.C. & Xu, T. Shaping cells and organs in Drosophila by opposing roles of fat body-secreted Collagen IV and perlecan. Dev Cell 21, 245–256 (2011).

46. Töpfer, U. et al. AdamTS proteases control basement membrane heterogeneity and organ shape in Drosophila. Cell Rep 43, 114399 (2024).

47. Serna-Morales, E. et al. Extracellular matrix assembly stress initiates Drosophila central nervous system morphogenesis. Dev Cell 58, 825–835.e826 (2023).

48. Morrissey, M.A. & Sherwood, D.R. An active role for basement membrane assembly and modification in tissue sculpting. J Cell Sci 128, 1661–1668 (2015).

49. Díaz-de-la-Loza, M.D. & Stramer, B.M. The extracellular matrix in tissue morphogenesis: No longer a backseat driver. Cells Dev 177, 203883 (2024).

50. Walther, R.F., Lancaster, C., Burden, J.J. & Pichaud, F. A dystroglycan-laminin-integrin axis coordinates cell shape remodeling in the developing Drosophila retina. PLoS Biol 22, e3002783 (2024).

51. Kimble, J. & Hirsh, D. The postembryonic cell lineages of the hermaphrodite and male gonads in Caenorhabditis elegans. Dev Biol 70, 396–417 (1979).

52. Barton, M.K., Schedl, T.B. & Kimble, J. Gain-of-function mutations of fem-3, a sex-determination gene in Caenorhabditis elegans. Genetics 115, 107–119 (1987).

53. Haag, E.S., Wang, S. & Kimble, J. Rapid Coevolution of the Nematode Sex-Determining Genes <EM>fem-3</EM> and <EM<tra-2</EM<. Current Biology 12, 2035–2041 (2002).

54. Kelley, C.A., et al. FLN-1/filamin is required to anchor the actomyosin cytoskeleton and for global organization of sub-cellular organelles in a contractile tissue. Cytoskeleton (Hoboken) 77, 379–398 (2020).

55. Merritt, C., Rasoloson, D., Ko, D. & Seydoux, G. 3’ UTRs are the primary regulators of gene expression in the C. elegans germline. Curr Biol 18, 1476–1482 (2008).

56. Zou, L., et al. Construction of a germline-specific RNAi tool in C. elegans. Scientific Reports 9, 2354 (2019).

57. Kovacevic, I. & Cram, E.J. FLN-1/filamin is required for maintenance of actin and exit of fertilized oocytes from the spermatheca in C. elegans. Dev Biol 347, 247–257 (2010).

58. Naba, A. Mechanisms of assembly and remodelling of the extracellular matrix. Nat Rev Mol Cell Biol 25, 865–885 (2024).

59. Haller, S.J., Roitberg, A.E. & Dudley, A.T. Steered molecular dynamic simulations reveal Marfan syndrome mutations disrupt fibrillin-1 cbEGF domain mechanosensitive calcium binding. Scientific Reports 10, 16844 (2020).

60. Chen, D.B. & Zheng, J. Regulation of placental angiogenesis. Microcirculation 21, 15–25 (2014).

61. Burton, G.J., Charnock-Jones, D.S. & Jauniaux, E. Regulation of vascular growth and function in the human placenta. REPRODUCTION 138, 895–902 (2009).

62. Orvik, A.B., et al. Plasma fibulin-1 levels during pregnancy and delivery: a longitudinal observational study. BMC Pregnancy Childbirth 21, 629 (2021).

63. Adam, S., et al. Binding of fibulin-1 to nidogen depends on its C-terminal globular domain and a specific array of calcium-binding epidermal growth factor-like (EG) modules. J Mol Biol 272, 226–236 (1997).

64. Hofmann, H., Voss, T., Kühn, K. & Engel, J. Localization of flexible sites in thread-like molecules from electron micrographs. Comparison of interstitial, basement membrane and intima collagens. J Mol Biol 172, 325–343 (1984).

65. Xu, X., et al. Age-related Impairment of Vascular Structure and Functions. Aging Dis 8, 590–610 (2017).

66. Shin, S.H., Lee, Y.H., Rho, N.-K. & Park, K.Y. Skin aging from mechanisms to interventions: focusing on dermal aging. Frontiers in Physiology 14 (2023).

67. Yasmin et al. The matrix proteins aggrecan and fibulin-1 play a key role in determining aortic stiffness. Scientific Reports 8, 8550 (2018).

68. Stiernagle, T. Maintenance of C. elegans. WormBook, 1–11 (2006).

69. Tuli, M.A., Daul, A. & Schedl, T. Caenorhabditis nomenclature. WormBook 2018, 1–14 (2018).

70. Dickinson, D.J. & Goldstein, B. CRISPR-Based Methods for Caenorhabditis elegans Genome Engineering. Genetics 202, 885–901 (2016).

71. Persson, K.E., Astermark, J., Björk, I. & Stenflo, J. Calcium binding to the first EGF-like module of human factor IX in a recombinant fragment containing residues 1-85. Mutations V46E and Q50E each manifest a negligible increase in calcium affinity. FEBS Lett 421, 100–104 (1998).

72. Sturm, Á., Saskoi, É., Tibor, K., Weinhardt, N. & Vellai, T. Highly efficient RNAi and Cas9-based auto-cloning systems for C. elegans research. Nucleic Acids Res 46, e105 (2018).

73. Schindelin, J., et al. Fiji: an open-source platform for biological-image analysis. Nature Methods 9, 676–682 (2012).

